# Proteomic analysis of human leptomeningeal matrisome identifies changes in Alzheimer’s disease

**DOI:** 10.64898/2026.05.27.728244

**Authors:** Kristy Urquhart, Alia Alia, Hui Chen, Lasanthi Jayathilaka, Cornelius Lam, Christopher Janson, Liudmila Romanova

## Abstract

The leptomeninges (arachnoid and pia) are the inner layers of the meninges that envelope the brain. They define the borders of the subarachnoid space (SAS) where cerebrospinal fluid (CSF) circulates. Leptomeninges act as the barrier tissue and contribute to fluid exchange and immune function of the brain. These processes are identified as key factors in the development of neurodegenerative diseases. The extracellular matrix (ECM) is a critical structural component of leptomeninges. However, it remains insufficiently characterized in health and disease. Here, we performed complete proteomic profiling of ECM proteins (matrisome) of human leptomeninges from individuals with Alzheimer’s disease (AD) and cognitively normal controls (CN) using sequential biochemical fractionation coupled with mass spectrometry. We resolved leptomeningeal matrisome based on the biochemical properties reflected by the protein solubility. This approach identified 211 ECM proteins in human leptomeninges and revealed disease-associated shifts in leptomeningeal solubility consistent with altered matrix organization and signaling. We found that ECM-linked regulators and secreted factors extracellular sulfatase SULF2, antithrombin-III (SERPINC1), secreted frizzled-related protein 3 (sFRP3) and integrin beta-5 (ITGB5), which is a receptor for fibronectin, were differentially expressed in AD leptomeninges based on the disease status. Seven ECM proteins were identified only in AD samples, and two ECM proteins in CN samples. Pathway enrichment implicated TGF-β and Wnt-related signaling and coagulation as critical for leptomeningeal function. Our findings provide the first comprehensive proteomic characterization of the human leptomeningeal matrisome and establish a biochemical framework for investigating how meningeal matrix remodeling contributes to AD.

## INTRODUCTION

Meninges are multi-layered membranes that envelop the brain and spinal cord and are essential to the structure and function of the central nervous system (CNS). While traditionally regarded as inert protective coverings, they are now recognized as highly specialized tissues with many roles and as dynamic participants in CNS physiology^1,2^. Meninges are vascularized, innervated, and immunologically active, serving as a critical interface between the CNS and peripheral systems^2–5^. Recent work has highlighted the diverse roles of the meninges in neurovascular regulation, immune surveillance, and tissue repair. Dysregulation of meningeal structure and function has been implicated in a range of neurological pathologies, including neurodegenerative diseases^6–8^, neuroinflammatory disorders^3,9^, and traumatic brain injury^10,11^, emphasizing their relevance to the CNS in health and disease.

Anatomically, meninges consist of three closely connected layers (Fig. 1). The dura mater is the outermost meningeal layer that attaches to the skull, providing stability and protection against mechanical force^10,12–14^. It is composed primarily of dense, collagen-rich connective tissue populated by fibroblasts and immune cells^12,14,15^. Dura is highly vascularized with blood and lymphatic vessels^16–18^. The arachnoid lies beneath the dura and forms the boundary of the subarachnoid space (SAS)^2,19^. The barrier functions of this layer are related to its epithelial-like structure, formed by flattened arachnoid barrier cells connected by tight junctions, creating a diffusion-limiting interface that regulates molecular exchange between the dura and cerebrospinal fluid (CSF)^2,20–22^. A basement membrane underlies the arachnoid barrier cell layer and is composed of extracellular matrix (ECM) proteins, including laminins, collagen IV, and nidogens, that contribute to mechanical stability ^23,24^. Arachnoid trabeculae extend from the inner arachnoid surface across the subarachnoid space to the pia mater, forming a structural framework that supports the brain and spinal cord within the CSF-filled compartment^2,19^.

**Figure 1.**
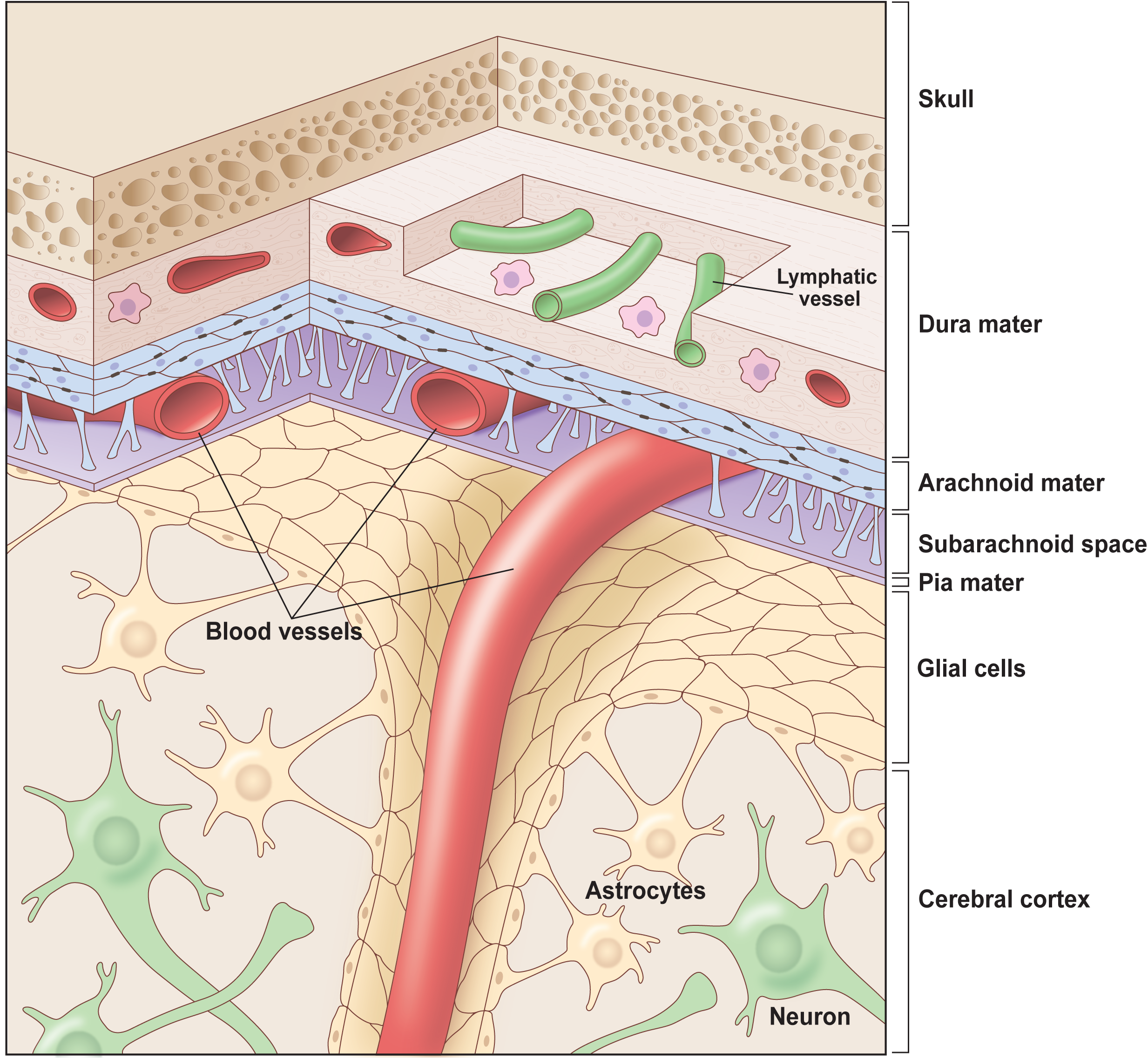
Schematic representation of meningeal layers. The outer layer of the dura is adjacent to the skull and is populated with fibroblasts and immune cells, has fenestrated blood vessels and lymphatic vessels. Arachnoid mater cells are connected with tight junctions and desmosomes and separate the cerebrospinal fluid (CSF) that circulates in the subarachnoid space (SAS) from the fenestrated dural vasculature. The pia is a thin, delicate layer that closely adheres to the brain surface, surrounds penetrating vessels, and interfaces with the glia limitans, thereby forming the innermost meningeal barrier between CSF and the brain parenchyma.

The pia mater, beneath the arachnoid, forms the innermost meningeal covering, a highly vascularized membrane that closely adheres to the brain and spinal cord. It contacts the glial limitans and creates a boundary between the cortex and SAS^5^. The pial basement membrane is essential for brain development. It anchors radial glia at the pial surface and provides the structural support neurons need to migrate to their correct position^25^. The arachnoid and pia are collectively called the leptomeninges, the layers of the meninges that are increasingly recognized for their roles in brain development and disease^26,27^.

Meningeal layers are rich in ECM, which includes proteins, polysaccharides, proteoglycans, and glycosaminoglycans (GAGs) that provide structural support and mediate cell-to-cell and cell-to-matrix interactions^28–30^. ECM proteins are collectively referred to as the matrisome^30,31^. Currently, the matrisome is classified into two groups: the core matrisome, which includes structural proteins such as collagens, proteoglycans, and glycoproteins, and the matrisome-associated proteins, which include enzymes and other factors that regulate matrix organization and remodeling^30–32^. The dura is composed primarily of the core ECM, dominated by collagen fibers that form the bulk of its structure^10,12,33^. By contrast, the ECM composition of the leptomeninges is much less understood. In the arachnoid layer, ECM proteins are found in SAS in trabeculae, the spiderweb-like collagen columns suspended in CSF, with parallel collagen bundles surrounded by fibroblasts^34^. In addition, ECM proteins, including laminins, collagen IV, nidogens, and heparan sulfate proteoglycans, contribute to basement membrane formation and bind cell-surface receptors such as integrins and dystroglycan, supporting the intercellular junctions that underlie arachnoid barrier function^20,24,25,29,35^. The pia mater is anchored by a highly specialized basement membrane enriched in laminins, nidogen, and collagen types IV, XV, and XVIII^23^. ECM-cell interactions at the pia-radial glia interface are required for normal cortical and cerebellar organization, as demonstrated by the requirement of β1-integrins for proper lamination^24^. Similarly, dystroglycan complexes at the pial surface regulate both neuronal and glial behaviors, highlighting the ECM’s dual structural and signaling roles^36^. The importance of ECM adhesion pathways is further shown by the essential role of α6-integrins in cortical and retinal lamination^35^. However, despite its structural and functional importance, this emerging understanding of early developmental roles, the meningeal matrisome, remains largely uncharacterized, particularly in adult human tissue and in the context of health and disease. Recent studies have begun to bridge this gap by defining ECM presence and abundance, confirming that ECM proteins constitute a substantial portion of the meningeal structure^1,27,33^. Transcriptomic analyses of mouse meninges revealed age-associated shifts in ECM gene expression, suggesting altered matrix production, turnover, and remodeling^3^. These transcriptional changes include pathways related to cell-matrix interactions, suggesting that aging may modify how meningeal cells interact with their surrounding extracellular environment. Recent proteomic and imaging studies further report increased collagen deposition near dura-associated lymphatic vessels and propose that this structural remodeling may influence lymphatic drainage^6^. However, the complete matrisome of each individual meningeal compartment remains to be fully characterized. Most previous studies examining ECM components in the meninges were performed in mice, where dura and leptomeninges cannot be physically separated and studied individually despite their distinct physiological roles.

Neuroimaging studies of cerebrovascular disorders demonstrate that alterations in ECM composition disrupt interstitial fluid movement and vascular integrity, implicating matrix remodeling as a contributor to Alzheimer’s disease (AD) pathology^37^. Experimental mouse AD models show that dysregulated hyaluronan remodeling worsens memory deficits, in part by interfering with clearance and autophagy pathways^38^. In cerebral amyloid angiopathy (CAA), abnormal deposition of matrix-associated proteins, fibrinogen, serum amyloid A, and apolipoprotein E, within leptomeningeal vessels promotes inflammation and vascular fragility^8^. Across neurodegenerative conditions, from AD to CAA, changes in matrix composition repeatedly emerge as drivers of pathology.

In this study, we, for the first time, characterized the matrisome of human leptomeninges and investigated its remodeling in AD based on the expression and solubility. Using a sequential protein-fractionation workflow, we extracted proteins across solubility classes, allowing comprehensive recovery of both soluble and insoluble components of the leptomeningeal matrix. This approach preserved key structural and regulatory elements of the ECM and enabled a high-resolution proteomic characterization of the leptomeninges. We identified 2926 total proteins, of which 211 are ECM proteins, and analyzed their differential expression between AD and cognitively normal human leptomeninges.

These findings provide the first in-depth proteomic atlas of the human leptomeningeal ECM. Our work provides a foundation for understanding how leptomeningeal ECM contributes to CNS health and disease. This dataset also creates new opportunities for biomarker discovery and therapeutic targeting in conditions where ECM dysregulation disrupts neurovascular, immune, and clearance pathways that drive neurodegeneration.

## METHODS

### Tissue procurement and characterization

Human leptomeningeal tissue was obtained from the Banner Sun Health Research Institute Brain and Body Donation Program (Banner Health, Sun City, AZ)^39^. Tissue samples were collected at autopsy under a rapid post-mortem protocol, with post-mortem intervals ranging from approximately 2 to 4 hours. Leptomeninges were carefully dissected from the cerebral surface during autopsy, snap-frozen, and stored at -80 °C until further processing. Specimens were selected based on comprehensive neuropathological assessment and clinical diagnostic data provided by the Brain and Body Donation Program^39^. Two groups were included in the samples utilized for this study: individuals with a neuropathologically confirmed diagnosis of AD (presence of cerebral amyloid angiopathy (CAA), Braak Stage V-VI, frequent neuritic plaques) and a cognitively normal (CN) control group (no CAA, Braak stage I-II-III, absent or minimal plaque pathology). A total of 10 cases were analyzed, comprising 5 AD cases (3 males, 2 females) and 5 CN cases (2 males, 3 females). All procedures involving human subjects were performed in accordance with the approved protocol of the Institutional Review Board of Banner Sun Health Research Institute Brain and Body Donation Program of Sun City, Arizona.

### Protein extraction and fractionation

Flash-frozen tissues were pulverized under liquid nitrogen. Pulverized tissue was suspended in extraction buffer containing 500 mM NaCl and 100 mM HEPES (pH 7.4). Samples were homogenized using a BeadBug homogenizer (Benchmark Scientific) with zirconium beads at maximum speed for two 15-second cycles. Homogenates were rotated for 2.5 hours, then centrifuged at 16,000 × g for 10 minutes at 23 °C. The pellet was washed with an equal volume of ice-cold ultra-pure water for 10 minutes. The pellet was then resuspended in SDS buffer containing 0.75% sodium deoxycholate and 0.3% SDS. Samples were vortexed and rotated overnight, followed by centrifugation at 16,000 × g for 10 minutes at 23 °C, and the supernatant was collected. The pellet was washed twice with ultra-pure water and centrifuged after each wash. The washed pellet was resuspended in 4 M guanidine hydrochloride (GuHCl) and 50 mM sodium acetate buffer. Samples were sonicated for 30 seconds at an amplitude level of 4 and incubated with gentle rotation for 72 hours. After incubation, the samples were centrifuged at 16,000 × g for 10 minutes at 23 °C and the supernatant was collected for LC-MS. The final pellet was washed three times in phosphate-buffered saline (PBS), then stored at −80 °C for subsequent mass spectrometry analysis. Protein concentration in each biochemical fraction was determined using the BCA assay (Pierce™ BCA Protein Assay Kit, Thermo Fisher Scientific) according to the manufacturer’s instructions.

### Sample preparation for Liquid Chromatography-Tandem Mass Spectrometry (LC-MS/MS)

All protein fractions were processed using a standardized tryptic digestion protocol. Protein digestion was performed using the S-Trap Micro kit (ProtiFi). Proteins were reduced with 10 mM dithiothreitol (DTT) at 37 °C for 90 minutes and alkylated with 25 mM iodoacetamide (IAA) for 30 minutes in a dark room at room temperature. Samples were acidified with phosphoric acid (12%) to a final concentration of 1.2%, then mixed with S-Trap binding buffer (100 mM triethylammonium bicarbonate [TEAB] in 90% methanol) and loaded onto S-Trap columns and centrifuged. Deglycosylation was performed on-column with PNGase F (New England Biolabs) at 37 °C for 2 hours in 0.1 M ammonium bicarbonate with continuous agitation. Sequential enzymatic digestion was then carried out: proteins were initially incubated with 1 μg Lys-C for 2 hours at 37°C, followed by overnight digestion with 3 μg trypsin (Promega). Peptides were eluted using sequential washes with TEAB, 0.2% formic acid (FA), and 50% acetonitrile/0.2% FA, combined and concentrated to dryness using a vacuum centrifuge (SpeedVac). Peptides were desalted using Oasis PRiME HLB cartridges (Waters), dried again, and reconstituted in 5% acetonitrile/0.1% formic acid for LC-MS/MS analysis. Final peptide concentrations were quantified, and aliquots were prepared for injection. Reduction and alkylation were performed as described above for the in-solution fractions, followed by dilution to 2 M urea and deglycosylation with PNGase F. Lys-C and trypsin digestion proceeded identically, and digests were centrifuged to remove debris. Supernatants were desalted via Oasis PRiME HLB cartridges and dried before LC-MS/MS.

### LC-MS/MS

Peptide samples were analyzed on a Q Exactive HF Hybrid Quadrupole-Orbitrap Mass Spectrometer coupled to an UltiMate 3000 RSLCnano System with a Nanospray Flex Ion Source (Thermo Fisher Scientific). Peptides were first loaded onto a Waters nanoEase M/Z C18 trap column (100 Å, 5 μm, 180 μm × 20 mm) and separated on an analytical Waters BEH C18 column (130 Å, 1.7 μm, 75 μm × 150 mm) at a flow rate of 300 nL/min. The mobile phases consisted of solvent A (0.1% formic acid in water) and solvent B (0.1% formic acid in 80% acetonitrile). A linear gradient was applied as follows: 5% B at 0–3 min, 8% B at 3.2 min, 8–30% B from 3.2–110 min, 35–95% B from 110–119 min, maintained at 95% B until 129 min, and re-equilibrated at 5% B until 140 min. Full MS scans were acquired in the Orbitrap over an m/z range of 350–1400 with a resolution of 120,000 (at m/z 200), an AGC target of 3 × 10⁶, and a maximum injection time of 50 ms. The top 20 most intense precursor ions with charge states two through five were selected for fragmentation by higher-energy collisional dissociation (HCD) at 28% normalized collision energy. Ions were temporarily excluded for 30 seconds if they fell within a 2.4 m/z range (±1.2 m/z) after being fragmented. MS/MS spectra were acquired with a resolution of 30,000, AGC target of 1 × 10⁵, and maximum injection time of 120 ms.

### Protein identification

Raw data files were analyzed using MaxQuant (v2.0.3.1) with the integrated Andromeda search engine. Spectra were searched against a curated Homo sapiens protein database (UniProt; reviewed and unreviewed entries), supplemented with a database of common laboratory contaminants. Search parameters included: parent ion mass tolerance of 10 ppm, fixed modification of carbamidomethylation on cysteine residues, and variable modifications including oxidation (M), deamidation (N/Q), hydroxylation (P/K), and N-terminal pyroglutamate formation. Peptide and protein identifications were filtered to achieve a 1% false discovery rate (FDR) at both peptide and protein levels. Only proteins identified by at least one unique peptide were retained for downstream analyses. Resulting identifications and quantitative data were imported and further processed using Scaffold DDA (v6.0.1; Proteome Software), for normalization, spectral counting, and exploratory analyses. Protein abundance was estimated semi-quantitatively based on the total number of spectral counts across samples. Principal component analysis (PCA) was performed on normalized spectral count data to assess sample clustering and potential batch effects.

### Data analysis and visualization

A protein was considered to be present if spectral counts were detected in at least three out of five biological replicates within a given experimental group. Statistical modeling of protein expression data was performed using the limma package in R^40^. Prior to modeling, protein abundance values were log2-transformed and normalized, log2 transformation was applied to approximate normality of spectral count data. Proteins were retained for analysis only if at least one fraction from one experimental group had no more than one sample with a missing value. Linear models were constructed with protein abundance modeled as a function of experimental covariates, group (AD vs. CN), and fraction. All statistical tests were two-sided. Empirical Bayes moderation of the standard errors was applied using the eBayes function to improve variance estimation in smaller sample sizes. P-values were corrected for multiple testing using the Benjamini-Hochberg method to control the false discovery rate^41^. Proteins were considered statistically significant if the FDR-adjusted p-value (q-value) was ≤ 0.05. Differential expression results were reported with corresponding log2 fold changes, raw p-values, and q-values. Visualization of differential expression data included volcano plots, dot/box plots, and heatmaps to highlight the most significantly regulated proteins and clustering patterns across samples. These visualizations were generated using the ggplot2 package in R^42^. Heatmaps were constructed to represent scaled protein intensities across groups and conditions, using hierarchical clustering to organize both samples and proteins. The protein data were annotated using MatrisomeAnalyzeR^31^. UniProt accession numbers were mapped to core matrisome and matrisome-associated categories based on the curated MatrisomeDB database^43^. These annotations stratified the proteomic findings by ECM subcategories, including collagens, glycoproteins, proteoglycans, ECM regulators, and affiliated proteins. Processed protein abundance tables exported from Scaffold DDA were curated and analyzed using tidyverse-based workflows (dplyr, tidyr, stringr), with additional analyses performed using limma, ggplot2, ComplexHeatmap, clusterProfiler, ReactomePA, and MatrisomeAnalyzeR^40,42,44,45^. Principal component analysis was performed in R using the prcomp() function on log2-transformed protein abundance values, and PCA plots were generated with ggplot2. Protein counts per fraction, and their proportional contribution to the total detected proteome, were calculated in R and visualized as bar charts and pie charts using GraphPad Prism (v9). Overlap of proteins between fractions was assessed using set-based comparisons in R, and fraction intersections were visualized using a four-way Venn diagram and an UpSet intersection plot generated with ggplot2-compatible tools.

### ECM-focused analyses

Proteins meeting the inclusion threshold (detected in ≥3 of 5 biological replicates per condition) were annotated using MatrisomeAnalyzeR to classify proteins into matrisome divisions (core matrisome, matrisome-associated, non-matrisome) and functional categories (collagens, ECM glycoproteins, proteoglycans, ECM regulators, ECM-affiliated proteins, and secreted factors) based on MatrisomeDB. Protein counts were summarized by biochemical fraction (NaCl, SDS, GuHCl, Pellet) and annotation class using R (v4.4.1) with tidyverse workflows. Distinct protein counts per fraction, division, and category were calculated using accession-level set operations. Absolute protein counts and fractional representations were visualized as stacked bar charts and pie charts to depict both total protein recovery and proportional ECM composition across fractions. All matrisome annotations, aggregation, and data structuring were performed in R, and final bar and pie chart visualizations were rendered using Adobe Illustrator (v29.8). To characterize changes in ECM protein solubility between AD and CN groups, a weighted solubility score was computed for each ECM protein in each experimental group. Sequential biochemical fractions were assigned an ordinal rank: NaCl = 1 (most soluble), SDS = 2, GuHCl = 3, and insoluble pellet = 4 (least soluble). For each protein, mean spectral count intensities were calculated per fraction per group, converted to within-group proportions totaling to 1.0 across fractions, and used as weights in a weighted mean of the fraction ranks. This yielded a single solubility score per protein per group, where a score approaching 1 indicates predominant detection in the NaCl fraction and a score approaching 4 indicates predominant detection in the insoluble pellet. The shift in solubility between groups was defined as the difference in solubility score (AD minus CN). Proteins were classified as shifting toward insolubility (positive difference) or toward solubility (negative difference). Shift magnitude was categorized using standard deviation-based thresholds derived from the empirical distribution of score changes across all 211 detected ECM proteins (standard deviation = 0.37): no shift (Δ score < 1 SD), low shift (1 SD ≤ Δ score < 2 SD), and high shift (Δ score ≥ 2 SD, i.e., ≥ 0.74). ECM proteins showing any directional shift were compiled into a supplementary table annotated by matrisome division, category, detected fractions in each group, and shift direction. A summary table reporting the dominant solubility trend per matrisome category was also generated. All analyses were performed in R using tidyverse workflows and the openxlsx package for table export.

### Data visualization

Differential abundance results from limma modeling were visualized using volcano plots, Venn diagrams, and summary plots that distinguished pairwise fraction-specific significance from group-level significance. Pairwise volcano plots were generated to display fraction-specific AD versus control differences, while a global volcano plot summarized proteins with significant Group Q-values across fractions. Venn diagrams were used to visualize the overlap of detected ECM proteins between AD and control samples. Functional pathway enrichment analyses (KEGG and Reactome) were performed on proteins meeting Group Q-value significance criteria. Enrichment results were visualized using bar plots and dot plots displaying gene ratios, protein counts, and FDR-adjusted q-values. All enrichment visualizations were generated using ggplot2 and exported as vector-based graphics. Hierarchical clustering heatmaps were generated to visualize protein distribution patterns across biochemical fractions and experimental groups. For these analyses, protein abundance values were normalized within samples and expressed as relative contribution across fractions. Proteins were annotated by ECM classification and category prior to visualization. Clustering was performed using hierarchical clustering with Euclidean distance and complete linkage, applied to both proteins and fractions. All figures were generated as publication-quality vector graphics using ggplot2 or ComplexHeatmap and exported as SVG or PDF files. Color schemes, legends, and axis labeling were standardized across figures to maintain visual consistency throughout the manuscript.

## RESULTS

### Sequential solubility-based extraction reveals complex proteomic profile in human leptomeninges

The goal of this study was to define composition of the matrisome in adult human leptomeninges and identify changes in the matrisome profile of individuals with AD. Human leptomeninges are a fragile and thin tissue with a shiny appearance that adheres to the surface of the postmortem brain after the dura mater is dissected (Fig. 2, A). Leptomeninges are largely avascular, but they contain discrete and prominent bridging vessels that traverse SAS and are heavily invested in arachnoid and pial tissues (Fig. 2, B). In contrast to rodent meninges, in which the dura and leptomeninges are tightly apposed and cannot be cleanly separated, human leptomeninges can be separated from the dura. This physical separation enables molecular analysis of ECM composition, specifically of leptomeningeal tissue without dura, which has high collagen content that can mask less abundant leptomeningeal ECM.

**Figure 2.**
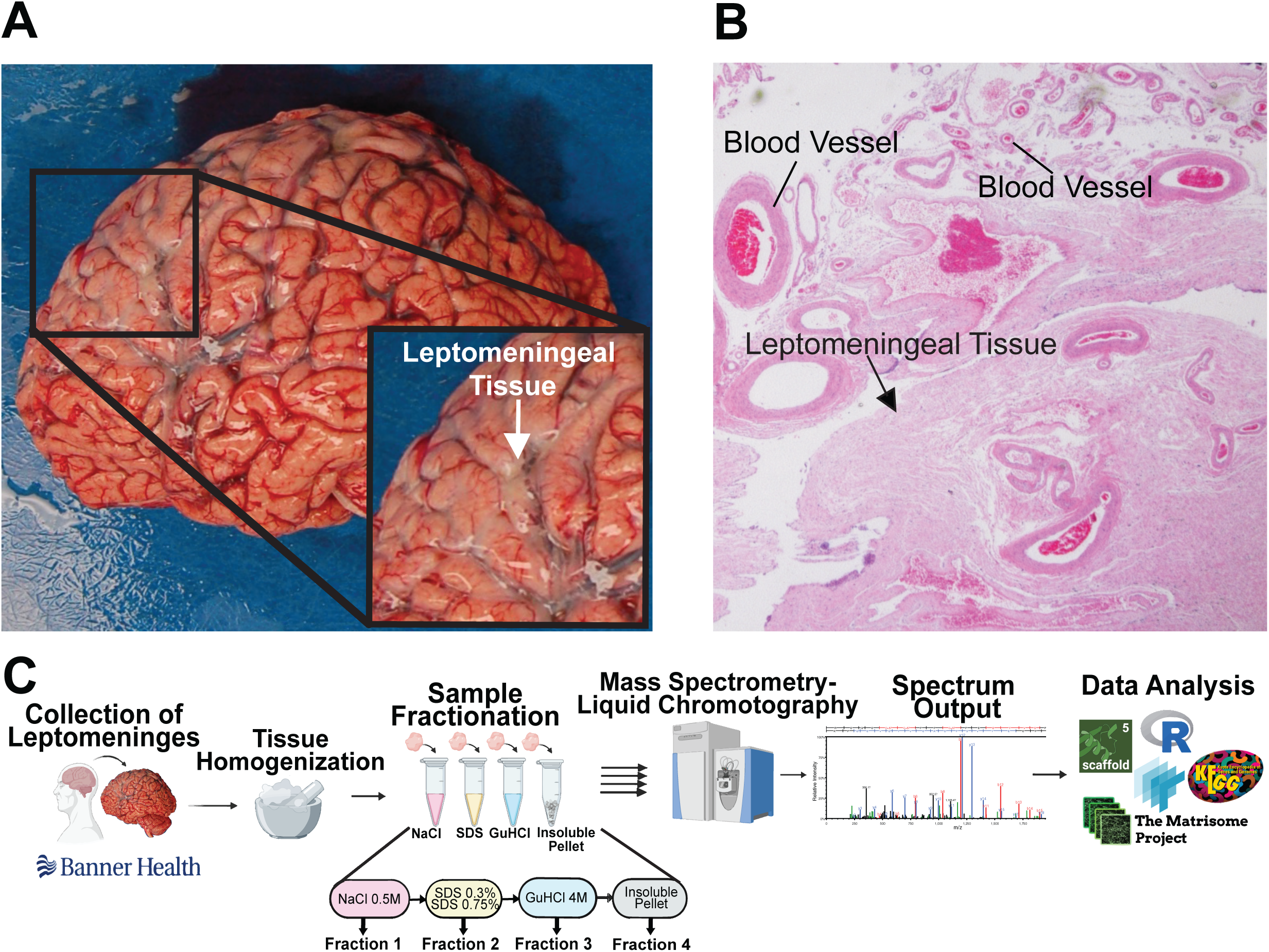
Human leptomeninges and the study workflow. (A) Human leptomeninges *in situ* on the cortical surface appear as a thin, translucent membrane. (B) Histological section of paraffin-embedded human leptomeninges showing vascular lumens surrounded by epithelial tissue and extracellular matrix. (C) Overview of the experimental workflow. Fresh-frozen leptomeninges were cryopulverized, homogenized, and were sequentially treated with the buffers of increasing detergent strength to solubilize proteins into distinct fractions. Each fraction was analyzed by mass spectrometry, and raw data were processed and analyzed as detailed in the Methods section. Images in (A) and (B) were provided by Banner Sun Health Research Institute.

We used well-characterized human leptomeningeal tissues provided by the Brain and Body Donation Program at Banner Sun Health Research Institute^39^. The samples were allocated into two groups. The cognitively normal (CN) group comprised five individuals aged 53–75 years (3 females, 2 males) with short post-mortem intervals ranging from 2.5-4.12 hours. Neuropathological evaluation confirmed the absence of AD pathology, with no CAA, zero plaque density, and low Braak stages (I–II). The AD group (AD) comprised five individuals aged 80–89 years (2 females, 3 males) with post-mortem intervals ranging from 2.08–3.33 hours. All AD specimens showed advanced neuropathology, including CAA, frequent amyloid plaques (12–15 total plaque counts, with frequent plaque density), and Braak stage VI neurofibrillary pathology. Detailed demographic and neuropathological information for all specimens is summarized in Table 1. The proteins were sequentially extracted and analyzed as outlined on Fig. 2, C. One of the potential challenges in specimen preparations for ECM analysis is incomplete extraction of ECM components, especially insoluble structural ECM proteins. In one-step extraction methods, less abundant proteins can be overpowered by a high structural ECM content^30,46^. To ensure complete ECM protein coverage, we used a stepwise optimized protein extraction with buffers of increasing strengths and concentration of solubilizing agents, and the inclusion of the insoluble final pellet for the proteomic analysis. We were able to achieve efficient homogenization without visible debris after bead-beating. The protocol produced sufficient protein quantities for mass spectrometry with the following fractions named based on the buffer composition: NaCl fraction, SDS fraction, GuHCl fraction, and the pellet. This approach permitted fractionation and analysis of the ECM proteins based on solubility, which is an important consideration because ECM remodeling is not defined solely by changes in total protein abundance, but also by shifts in biochemical accessibility, solubility, and matrix organization. These features were captured with a sequential, solubility-focused extraction workflow. For the data analysis, a protein was considered to be present if the spectral counts were detected in three out of five biological specimens within a given experimental group. This threshold was selected to increase sensitivity and capture of ECM proteins that, while less consistently detected, may nonetheless contribute biologically relevant information in the context of this study using human tissue with inherently high variability.

**Table 1.**
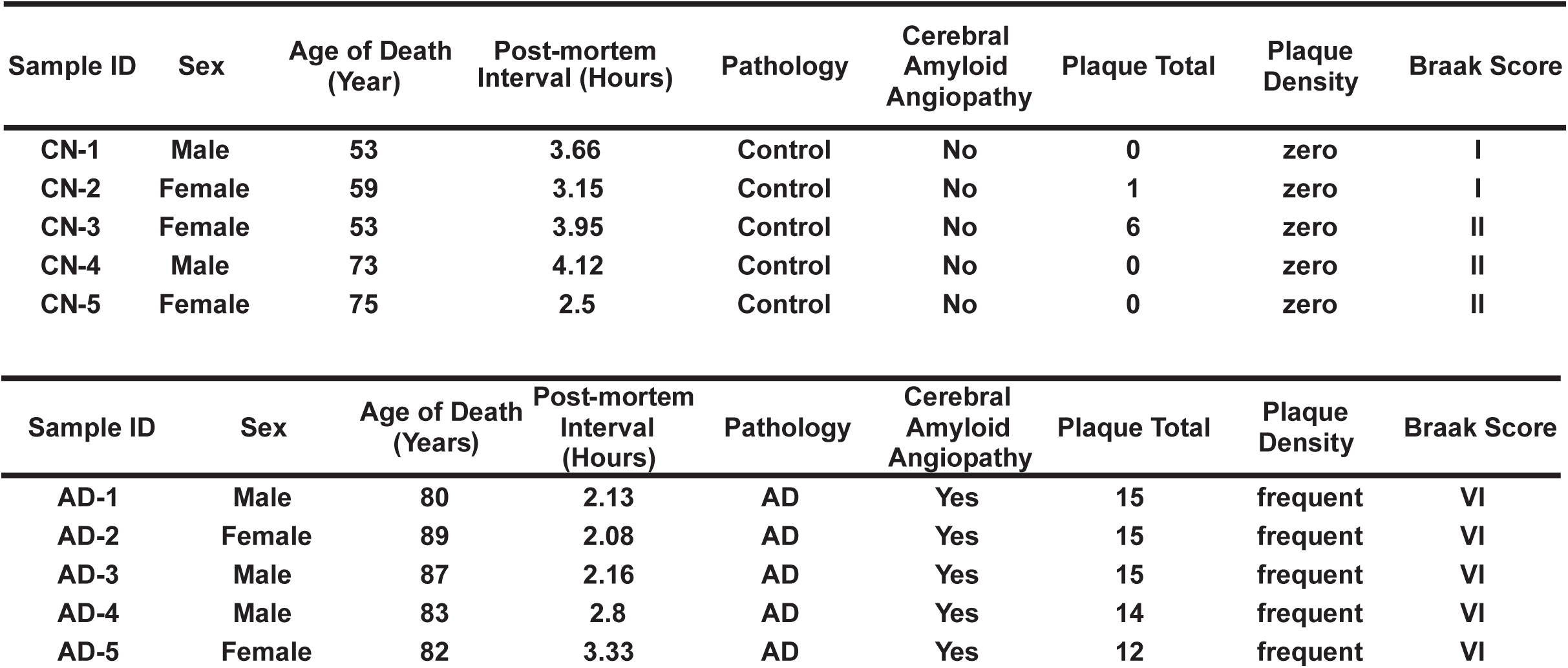
Demographic and neuropathological characteristics of human leptomeningeal specimens. Summary of donor sex, age at death, PMI, neuropathological diagnoses, CAA status, total and density of neuritic plaques, and Braak stage for all CN and AD specimens used in this study. Ages ranged from 53 to 75 years of age for CN specimens and 80 to 89 for AD. CAA was present in all AD samples and absent in all CN samples. All CN samples exhibited zero plaque burden and Braak stages I-II, while AD cases exhibited significant plaque burden (frequent) and Braak stage VI.

Principal component analysis showed that samples grouped predominantly by fraction rather than by the disease status, indicating that biochemical solubility exerts the strongest influence on proteomic variance in leptomeningeal tissue. Replicates clustered tightly, demonstrating strong technical reproducibility. NaCl, SDS, GuHCl, and pellet groups occupied distinct regions along PC1 and PC2, which accounts for 68.5% of total variance on Fig. 3, A. This fraction-driven separation stresses the need for a multi-fraction workflow to capture the biochemical diversity of the leptomeningeal proteome. Using this approach, we identified a total of 2926 proteins across all samples: 1523 in the NaCl fraction, 2600 in the SDS fraction, 564 in the GuHCl fraction, and 281 in the pellet (Fig. 3, B). The majority of the proteins were detected in SDS fraction with the least number of proteins in the pellet. The full list of all identified proteins can be found in the Supplementary Table 1. The proportional contribution of each fraction is further summarized in Fig. 3, C, where proteins in the SDS fraction account for 52% of all identifications, emphasizing the dominance of the SDS-soluble proteome. The SDS extraction step is typically enriched for intracellular and membrane-associated proteins. NaCl accounted for 31% of the identified proteins. This fraction primarily contains secreted and plasma proteins. ECM-enriched GuHCl and the pellet fractions accounted for 11% and 6%. Most of the proteins were identified in more than one fraction. Pie chart on Fig. 3, D and upset plot on Fig. 3, E show distribution of the identified proteins across the fractions. A detailed examination of these intersections revealed that NaCl and SDS have the most extensive unique protein subsets (265 and 1,238 proteins, respectively) showing effectiveness of SDS-based one-step extraction protocols. However, 31 proteins were uniquely identified in GuHCl and 4 in the pellet, showing that those proteins are mostly inaccessible with less aggressive extraction methods. These unique subsets highlight the contribution of low-solubility and structurally embedded proteins, further validating the importance of sequential extraction for comprehensive characterization of ECM. A shared core of 181 proteins was recovered in all four fractions. Together, these analyses demonstrated that the human leptomeningeal proteome is distributed across biochemically distinct fractions, with the highly insoluble components preferentially enriched in detergent-resistant chaotropic extracts.

**Figure 3.**
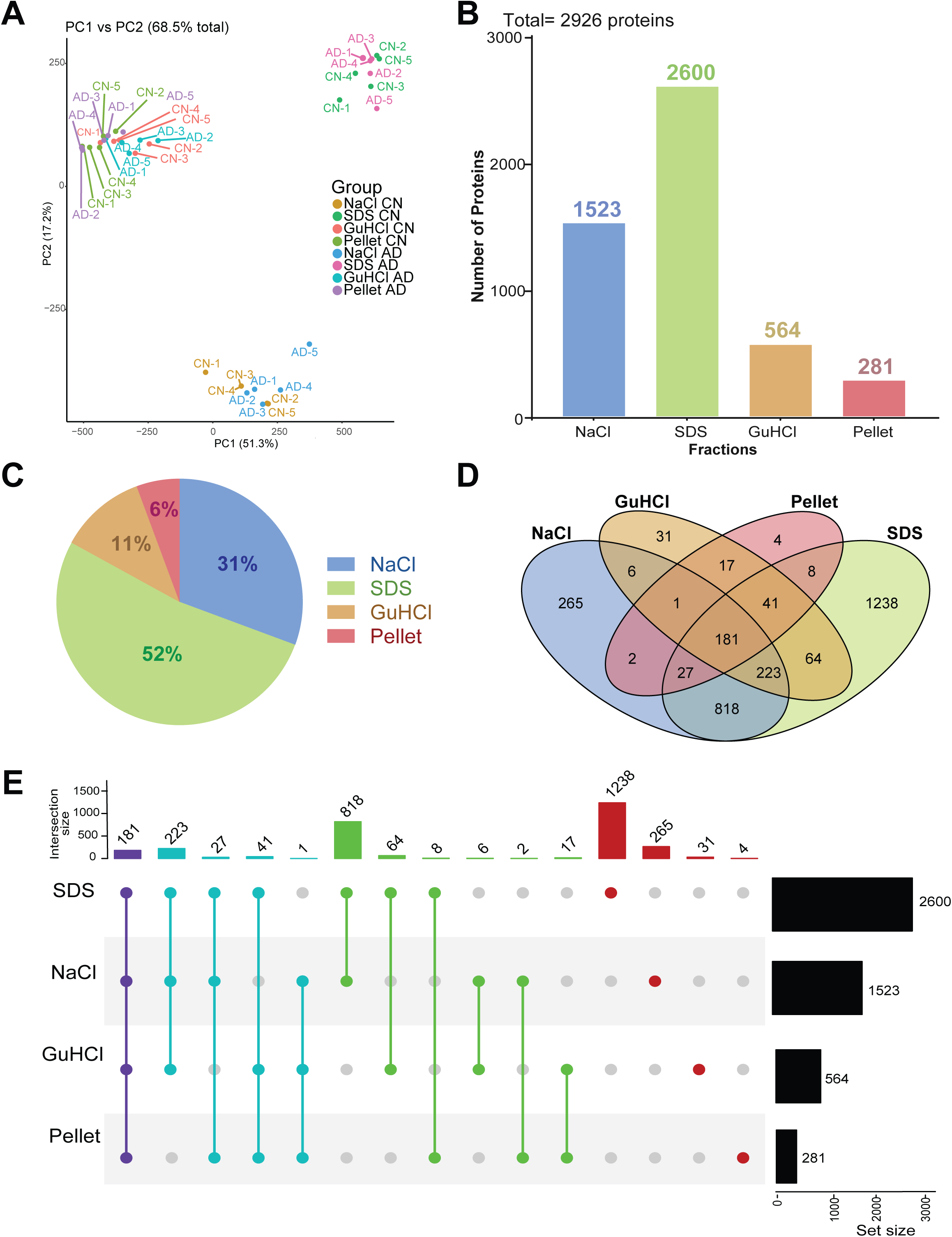
Distribution of the proteins identified in human leptomeninges across fractions. (A) Principal component analysis (PCA) plot of proteomic profiles across all samples, with each point representing a biological specimen. Proximity along the axes indicates similar expression patterns, while separation reflects distinct proteomic signatures. The first two principal components explain 51.3% (PC1) and 17.2% (PC2) of total variance of 68.5%. The samples cluster based on solubility rather than on the experimental group. (B) The graph shows the total number of proteins identified in each fraction. (C) Pie chart shows percentage of the total protein number identified in each fraction. (D) Venn diagram shows distribution of proteins across fractions. (E) Upset plot shows proteins across all experimental groups, with shared and unique proteins indicated on the plot. Proteins identified in all four fractions are shown in purple; in three out of four fractions are shown in blue; in two out of four fractions are shown in green. The proteins identified only in one fraction are shown in red.

### Characterization of human leptomeningeal matrisome

Identified proteins were annotated using MatrisomeDB, which functionally divides the ECM proteins into core and associated matrisome^43^. The core matrisome includes collagens, glycoproteins, and proteoglycans, and the associated matrisome includes regulators, ECM-affiliated proteins, and secreted factors. A total of 211 ECM proteins were identified across all fractions in both sample groups. Full list of ECM proteins, with accession numbers and divided by category, can be found in Supplementary Table 2. Fig. 4, A shows the breakdown of identified proteins by the matrisome type and ECM category in each fraction. The majority of non-matrisome proteins were extracted in the soluble fractions, NaCl, and SDS. The number of non-matrisome proteins decreases substantially in GuCl and the pellet fractions with the highest percentage of less soluble core-matrisome proteins in the pellet. The proportion of ECM-regulators, secreted factors, and proteoglycans remained roughly similar across fractions, but the collagens were mostly represented in the pellet. The protein counts (Fig. 4, B and C) and percentages (Fig. 4, D) of ECM-proteins by category and by fraction show that ECM proteins represent, overall, a smaller fraction of the total proteins extracted, highlighting the need for the fractionation approach. Glycoproteins represent the most abundant type of ECM-proteins in human leptomeninges with proteoglycans and secreted factors being least abundant (Fig. 4, E-G).

**Figure 4.**
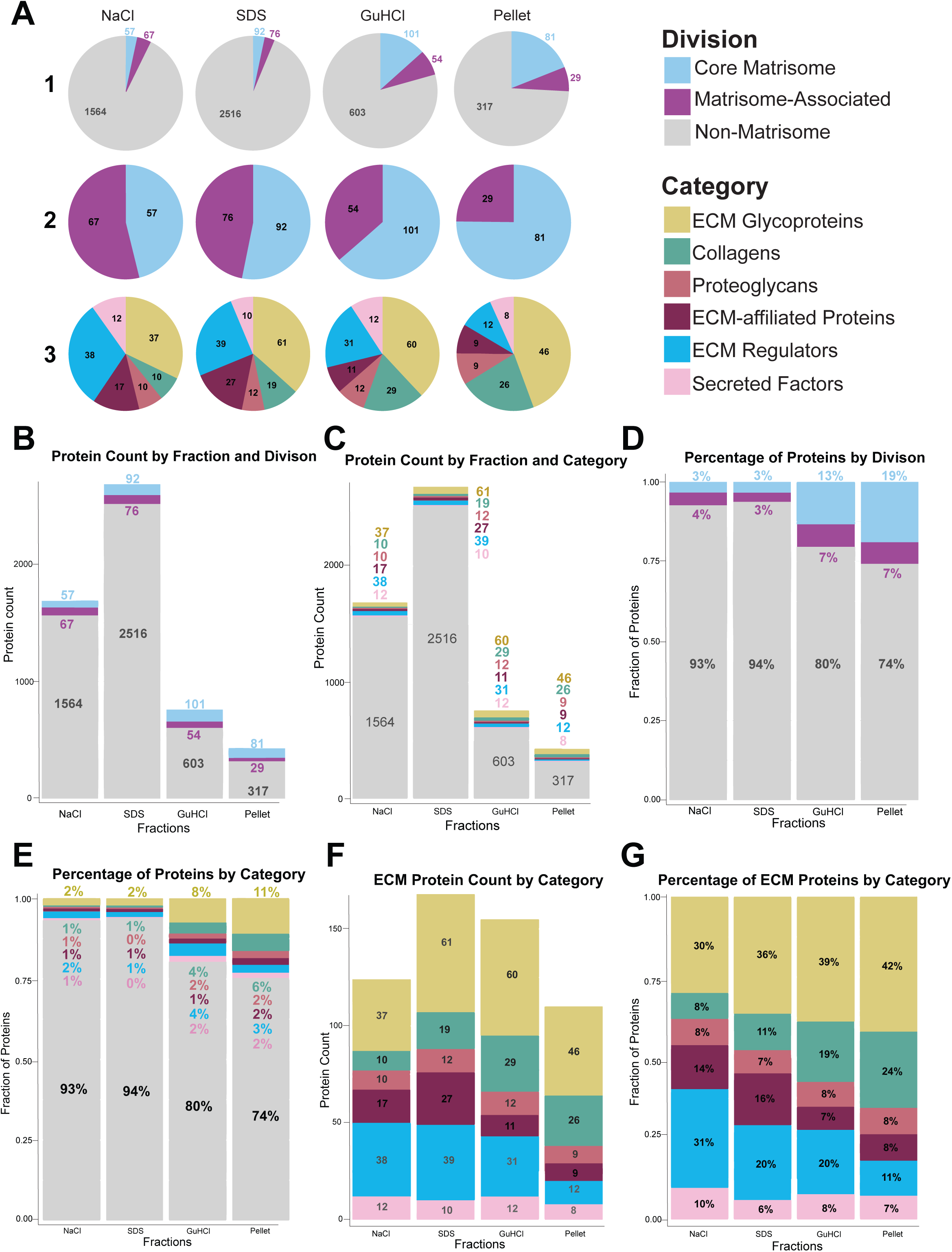
Sequential extraction reveals solubility-dependent organization of the human leptomeningeal matrisome. (A) Pie charts show the distribution of proteins across sequential extraction fractions (NaCl, SDS, GuHCl, and insoluble pellet) by division (core matrisome, matrisome-associated, non-matrisome) and by matrisome category (ECM glycoproteins, collagens, proteoglycans, ECM-affiliated proteins, ECM regulators, and secreted factors). Non-matrisome proteins comprise the majority of proteins in NaCl and SDS fractions, whereas the GuHCl and pellet fractions show increased representation of core matrisome components. (B) Bar graph of total protein counts per fraction by division, demonstrating progressive enrichment of core matrisome proteins in the less soluble GuHCl and pellet fractions. (C) Protein counts per fraction categorized into ECM glycoproteins, collagens, proteoglycans, ECM-affiliated proteins, ECM regulators, and secreted factors. (D) Distribution of proteins by division across extraction steps demonstrates increased proportion of core matrisome in less soluble fractions. (E) Distribution of proteins across fractions by category shows relative enrichment of ECM glycoproteins and collagens in the GuHCl and pellet fractions compared to more soluble fractions. (F) Absolute counts of ECM proteins per category in each fraction. (G) Percent of ECM proteins by category within each fraction.

We identified 29 collagens including chains of COL1; COL2A1, COL3A1; chains of COL4, COL5, COL6, COL8; COL10A1, COL11A2, COL12A1, COL21A1, COL16A1, COL15A1, COL27A1, COL18A1 and COL14A1. The proportion of the glycoproteins remained similar across the fractions. The glycoprotein group was primarily composed of basement membrane, elastic fiber, and perivascular ECM proteins. Elastic fiber and microfibril-associated proteins included elastin (ELN), fibrillin-1 (FBN1), fibulin family members (FBLN1, FBLN2, FBLN5), and microfibril-associated glycoproteins (MFAP2, MFAP4), and members of the EMILIN family (EMILIN1–3), which link elastic fibers to the surrounding ECM and regulate vascular function^47,48^. Multiple laminins were identified, including laminin α (LAMA1–5), β (LAMB1–2), and γ (LAMC1, LAMC3) subunits, together with nidogen-1 (NID1), nidogen-2 (NID2), and agrin (AGRN). Other perivascular ECM-associated proteins were nephronectin (NPNT), tubulointerstitial nephritis antigen–like protein (TINAGL1), extracellular matrix protein 2 (ECM2), and hemicentin-1 (HMCN1). Multiple ECM regulators associated with fibrotic remodeling and transforming growth factor–β (TGFβ) signaling were identified within the leptomeningeal matrisome. These included latent TGFβ-binding proteins (LTBP1, LTBP2, LTBP4), TGFβ-induced protein (TGFBI), periostin (POSTN), adipocyte enhancer-binding protein 1 (AEBP1), procollagen C-endopeptidase enhancer 1 (PCOLCE), and collagen triple helix repeat-containing protein 1 (CTHRC1). The leptomeningeal matrisome included proteins typically associated with vascular endothelium, such as von Willebrand factor (VWF), von Willebrand factor A domain–containing proteins (VWA1, VWA5A), multimerin-1 and -2 (MMRN1, MMRN2), matrix Gla protein (MGP), endothelial-derived IGFBP7, and EGF-like repeat–containing protein EDIL3^49–51^.

### ECM and non-ECM protein changes in Alzheimer’s disease

Human specimens for this study were selected to represent homogeneous sample groups based on age and pathological hallmarks. However, human tissues have inherent variability. To assess variability in ECM protein detection across the specimens, we separated the proteins into two groups based on the detection in three or more samples within each group and extraction fraction. Venn diagram on Fig. 5, A shows that under this criterion, 202 ECM proteins were consistently identified in both AD and CN leptomeninges in three or more cases. Seven ECM proteins were identified only in AD samples, and two ECM proteins met this threshold in CN samples (Fig. 5, B). The proteins identified exclusively in 3 or more specimens in AD group included EGF-like repeat and discoidin I-like domain-containing protein 3, Netrin-1, Matrix Gla protein, Midkine, Thrombospondin-2, extracellular sulfatase-2 (SULF2), and Cathepsin-F. The two ECM proteins meeting the threshold in CN samples were latent transforming growth factor beta-binding protein 4 (LTBP4) and Tenascin-R.

**Figure 5.**
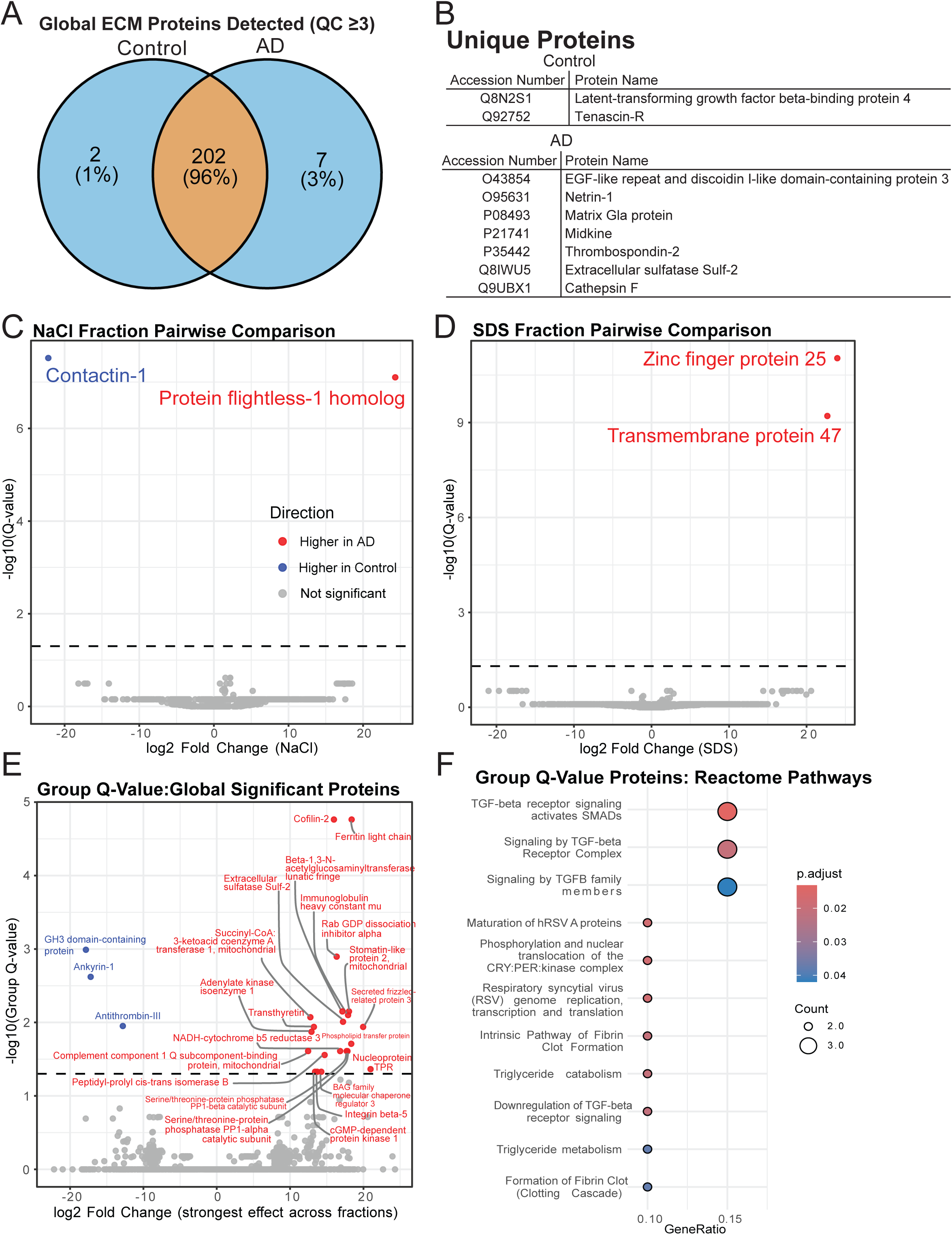
Pathway enrichment in human leptomeninges and changes in ECM and non-ECM protein expression and in Alzheimer’s disease. (A) A Venn diagram shows the proportion of unique and shared ECM proteins detected in three or more cases from CN and AD groups. (B) Proteins uniquely detected in AD or CN samples under the same criterion. (C) A volcano plot demonstrating pairwise comparison between proteins detected in the NaCl fraction between AD and CN samples. Points represent individual proteins plotted by log₂ fold change (AD vs control) and −log₁₀(Q-value). Proteins significantly enriched in AD are shown in red, those enriched in CN are in blue, and non-significant proteins are in gray. (D) A volcano plot showing pairwise comparison of proteins detected in the SDS fraction. (E) Differential protein analysis across all fractions based on the group, AD vs CN. Proteins are plotted according to their strongest observed effect across extraction fractions. The dashed horizontal line indicates the significance threshold based on group-level Q-values. Proteins downregulated in AD are shown in blue and upregulated in AD are shown in red. (F) Reactome pathway enrichment analysis of differentially expressed proteins. Enriched pathways include TGF-β receptor signaling, clotting cascade, intrinsic pathway of fibrin clot formation, and triglyceride metabolism and catabolism. Adjusted p-value indicated by the spot color. Gene count is reflected by the spot size.

The proteomic data was analyzed with pairwise differential analysis using two approaches. Most proteins were detected in multiple fractions. We first identified significant proteins within each biochemical fraction. As shown on the volcano plot on Fig. 5, C, two proteins showed statistically significant difference between AD and CN samples in NaCl fraction after multiple-comparison correction. Protein flightless-1 homolog (FLII) showed significantly higher abundance in the AD group, whereas Contactin-1 was significantly enriched in the CN group. In the SDS-soluble fraction, we identified statistically significant differences in the abundance of Zinc finger protein 25 (ZNF25) and Transmembrane protein 47 (TMEM47) levels, which were higher in AD compared to CN leptomeninges (Fig. 5, D). No proteins were found to be statistically different between GuHCl or the pellet fractions between AD and CN groups. This pairwise comparison between individual fractions showed that the disease effect was the strongest on the solubility of these proteins. Next, we analyzed the overall group effect (AD vs. CN) using the group Q-value from a multifactor generalized linear model to determine the disease-related effect independent of biochemical solubility. We identified 24 proteins, which were differentially expressed between AD and CN groups (Fig. 5, E). The list of the proteins with the accession numbers and functions is shown in Table 2 where ECM proteins are highlighted in green. The majority of the proteins were significantly enriched in AD, and a smaller set of proteins, such as antithrombin-III, ankyrin-1, and a protein with a GH3 domain, showed downregulation in AD compared to CN. Among these proteins, three proteins were identified as matrisome-associated: two ECM-regulators, extracellular sulfatase Sulf-2 (SULF2) and antithrombin-III (SERPINC1), and one secreted factor, secreted frizzled-related protein 3 (sFRP3). We also identified integrin beta-5 (ITGB5), which is not annotated as an ECM protein in MatrisomeDB but is closely related to ECM by mediating cell adhesion to extracellular matrix ligands^52^.

**Table 2.**
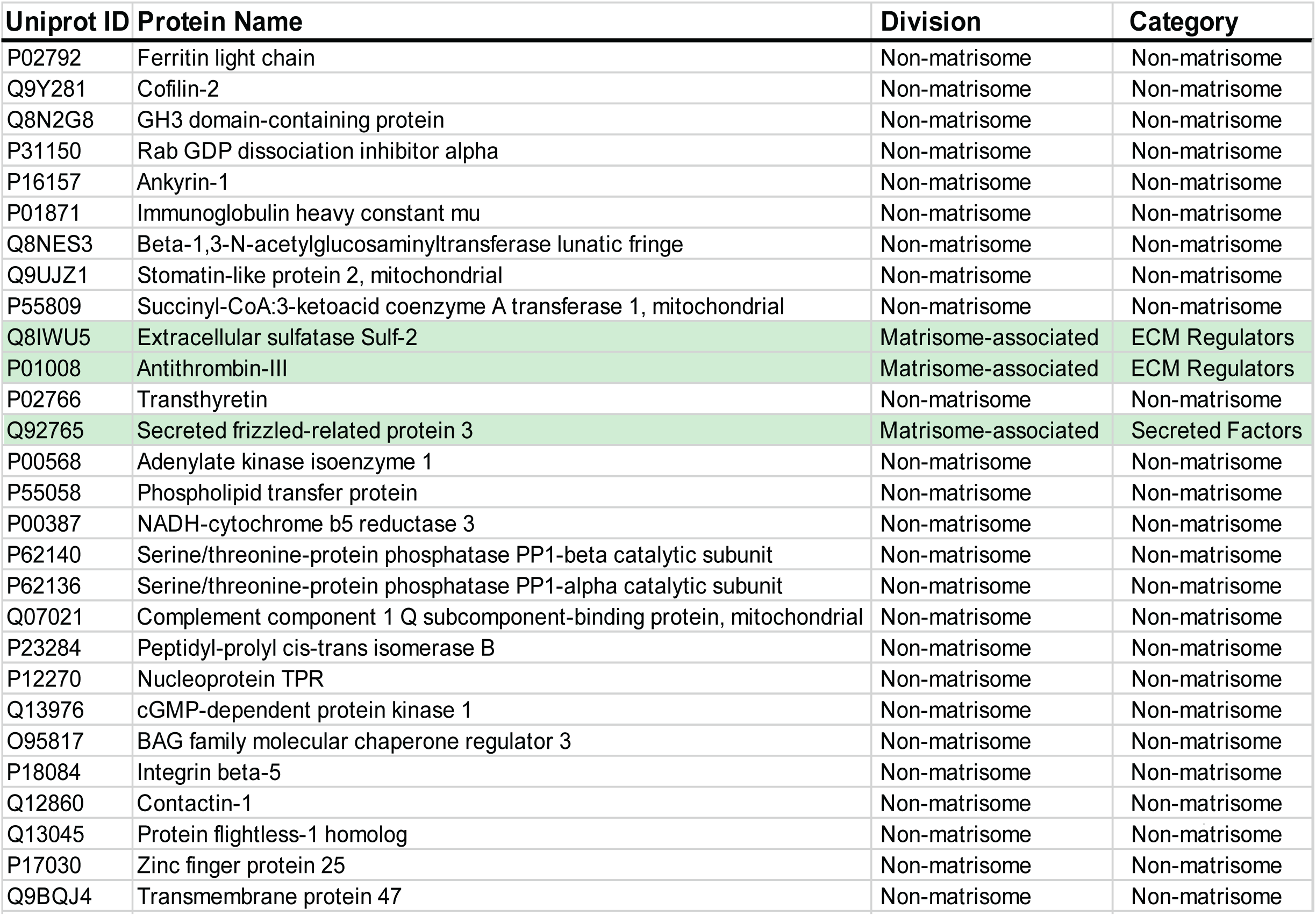
Statistically significant proteins between CN and AD sample groups. The proteins are organized by the division (core matrisome, associated matrisome and integrins) and by ECM type (collagens, glycoproteins, proteoglycans, secreted factors, ECM regulators and ECM-affiliated proteins). UniProt ID is displayed on the left from the protein name.

| Uniprot ID | Protein Name | Division | Category |
| --- | --- | --- | --- |
| P02792 | Ferritin light chain | Non-matrisome | Non-matrisome |
| Q9Y281 | Cofilin-2 | Non-matrisome | Non-matrisome |
| Q8N2G8 | GH3 domain-containing protein | Non-matrisome | Non-matrisome |
| P31150 | Rab GDP dissociation inhibitor alpha | Non-matrisome | Non-matrisome |
| P16157 | Ankyrin-1 | Non-matrisome | Non-matrisome |
| P01871 | Immunoglobulin heavy constant mu | Non-matrisome | Non-matrisome |
| Q8NES3 | Beta-1,3-N-acetylglucosaminyltransferase lunatic fringe | Non-matrisome | Non-matrisome |
| Q9UJZ1 | Stomatin-like protein 2, mitochondrial | Non-matrisome | Non-matrisome |
| P55809 | Succinyl-CoA:3-ketoacid coenzyme A transferase 1, mitochondrial | Non-matrisome | Non-matrisome |
| Q8IWU5 | Extracellular sulfatase Sulf-2 | Matrisome-associated | ECM Regulators |
| P01008 | Antithrombin-III | Matrisome-associated | ECM Regulators |
| P02766 | Transthyretin | Non-matrisome | Non-matrisome |
| Q92765 | Secreted frizzled-related protein 3 | Matrisome-associated | Secreted Factors |
| P00568 | Adenylate kinase isoenzyme 1 | Non-matrisome | Non-matrisome |
| P55058 | Phospholipid transfer protein | Non-matrisome | Non-matrisome |
| P00387 | NADH-cytochrome b5 reductase 3 | Non-matrisome | Non-matrisome |
| P62140 | Serine/threonine-protein phosphatase PP1-beta catalytic subunit | Non-matrisome | Non-matrisome |
| P62136 | Serine/threonine-protein phosphatase PP1-alpha catalytic subunit | Non-matrisome | Non-matrisome |
| Q07021 | Complement component 1 Q subcomponent-binding protein, mitochondrial | Non-matrisome | Non-matrisome |
| P23284 | Peptidyl-prolyl cis-trans isomerase B | Non-matrisome | Non-matrisome |
| P12270 | Nucleoprotein TPR | Non-matrisome | Non-matrisome |
| Q13976 | cGMP-dependent protein kinase 1 | Non-matrisome | Non-matrisome |
| O95817 | BAG family molecular chaperone regulator 3 | Non-matrisome | Non-matrisome |
| P18084 | Integrin beta-5 | Non-matrisome | Non-matrisome |
| Q12860 | Contactin-1 | Non-matrisome | Non-matrisome |
| Q13045 | Protein flightless-1 homolog | Non-matrisome | Non-matrisome |
| P17030 | Zinc finger protein 25 | Non-matrisome | Non-matrisome |
| Q9BQJ4 | Transmembrane protein 47 | Non-matrisome | Non-matrisome |

Several non-ECM proteins were differentially expressed in AD leptomeninges. Among proteins with significantly higher abundance in AD, protein flightless-1 homolog (FLII) is an actin-binding nuclear receptor coactivator^53^, and cofilin-2 is a regulator of actin filament dynamics^54^. Stomatin-like protein 2 is a mitochondrial membrane protein involved in mitochondrial biogenesis^55^, and succinyl-CoA:3-ketoacid coenzyme A transferase 1 is a mitochondrial enzyme central to ketone body metabolism^56^. Ferritin light chain is a component of the iron storage complex^57^, and BAG3 is a chaperone scaffolding protein that links HSP70 to the cellular stress response^58^. Elevated immunoglobulin heavy constant mu and complement C1q-binding protein were also identified in AD. Additional proteins upregulated in AD included lunatic fringe, a glycosyltransferase that modulates Notch receptor signaling^59^, zinc finger protein 25, a transcription factor^60^, and transmembrane protein 47, a member of the claudin superfamily involved in epithelial junction maturation^61^. Among proteins with lower abundance in AD, ankyrin-1 is a cytoskeletal scaffolding protein^62^, contactin-1 is a neuronal cell adhesion molecule involved in axogenesis and myelination^63^, and adenylate kinase isoenzyme 1 is an enzyme that maintains cellular adenine nucleotide balance^64^. Reduced PP1 phosphatase subunits PPP1CA and PPP1CB were also identified, along with cGMP-dependent protein kinase 1 and a GH3 domain-containing protein.

Reactome pathway analysis was conducted for statistically significant proteins. Enrichment highlighted a cohesive group of pathways related to TGF-β receptor signaling, fibrin clot formation, lipid metabolism, and cytoskeletal regulation (Fig. 5, F). The most significantly enriched pathways were associated with TGF-β signaling, including TGF-β receptor signaling activates SMADs, signaling by the TGF-β receptor complex, and signaling by TGF-β family proteins^44^. These enrichments were driven primarily by shared molecular nodes involving PPP1CA, PPP1CB, and ITGB5, reflecting enrichment of receptor-proximal phosphorylation and dephosphorylation processes rather than discrete downstream transcriptional programs^65–67^. Coagulation-associated pathways, including the Intrinsic pathway of fibrin clot formation and the formation of fibrin clot (clotting cascade), were significantly enriched, mapping to proteins such as SERPINC1 and C1QBP.

### Alzheimer’s disease is associated with solubility-dependent redistribution of the leptomeningeal matrisome

Hierarchical clustering of protein abundance expressed as percent of total signal per protein across fractions segregated samples primarily by extraction fraction rather than diagnosis, confirming strong biochemical partitioning of leptomeningeal ECM components (Fig. 6). However, within each fraction, AD samples exhibited distinct redistribution patterns relative to controls (Supplementary Table 3). We identified 73 matrisome proteins with detectable solubility shifts in AD relative to controls, 38 exhibited a shift toward increased biochemical insolubility, while 35 showed a shift toward greater solubility, with increased representation in NaCl or SDS fractions in AD (Table 3). Among proteins shifting toward insolubility, ECM glycoproteins were prominently represented, including papilin (PAPLN), ECM2, multimerin-1 (MMRN1), cochlin (COCH), EDIL3, netrin-1 (NTN1), hemicentin-1 (HMCN1), IGFBP7, and both EMILIN-1 and EMILIN-3. Collagens shifting toward insolubility included COL4A4 and COL15A1, as well as COL27A1. Aggrecan (ACAN) was the sole proteoglycan exhibiting increased insolubility. Among ECM regulators, multiple cathepsins (CTSH, CTSL, CTSG), serpins (SERPINA3, SERPINH1), inter-alpha-trypsin inhibitor heavy chain H1 (ITIH1), ADAM10, cystatin-C (CST3), and serpin H1 (SERPINH1) all shifted toward less soluble fractions. Secreted factors with increased insolubility included several S100 family members (S100B, S100A11, S100A9, S100A1), midkine (MDK), pleiotrophin (PTN), and complement C1q subunit B (C1QB). Several ECM proteins showed increased solubility in AD. All four laminin subunits, LAMA1, LAMA3, LAMA4, and LAMC3, appeared in more soluble fractions in AD relative to CN samples. Collagens COL8A1 and COL16A1 similarly gained representation in the NaCl fraction in AD. Additional ECM glycoproteins shifting toward solubility included elastin (ELN), periostin (POSTN), thrombospondin-2 (THBS2), LTBP1, nephronectin (NPNT), SPARCL1, SRPX, and SRPX2. Among ECM regulators, antithrombin-III (SERPINC1), TIMP3, serine protease 1 (PRSS1), and cathepsins CTSA, CTSF, and CTSC all shifted toward more soluble fractions. Secreted factors, including platelet basic protein (PPBP), interleukin-17D (IL17D), and Wnt-2b (WNT2B), also showed solubility increases in AD. Overall, solubility changes point to AD-associated remodeling of the leptomeningeal matrisome across structural, regulatory, and signaling components of the matrix.

**Figure 6.**
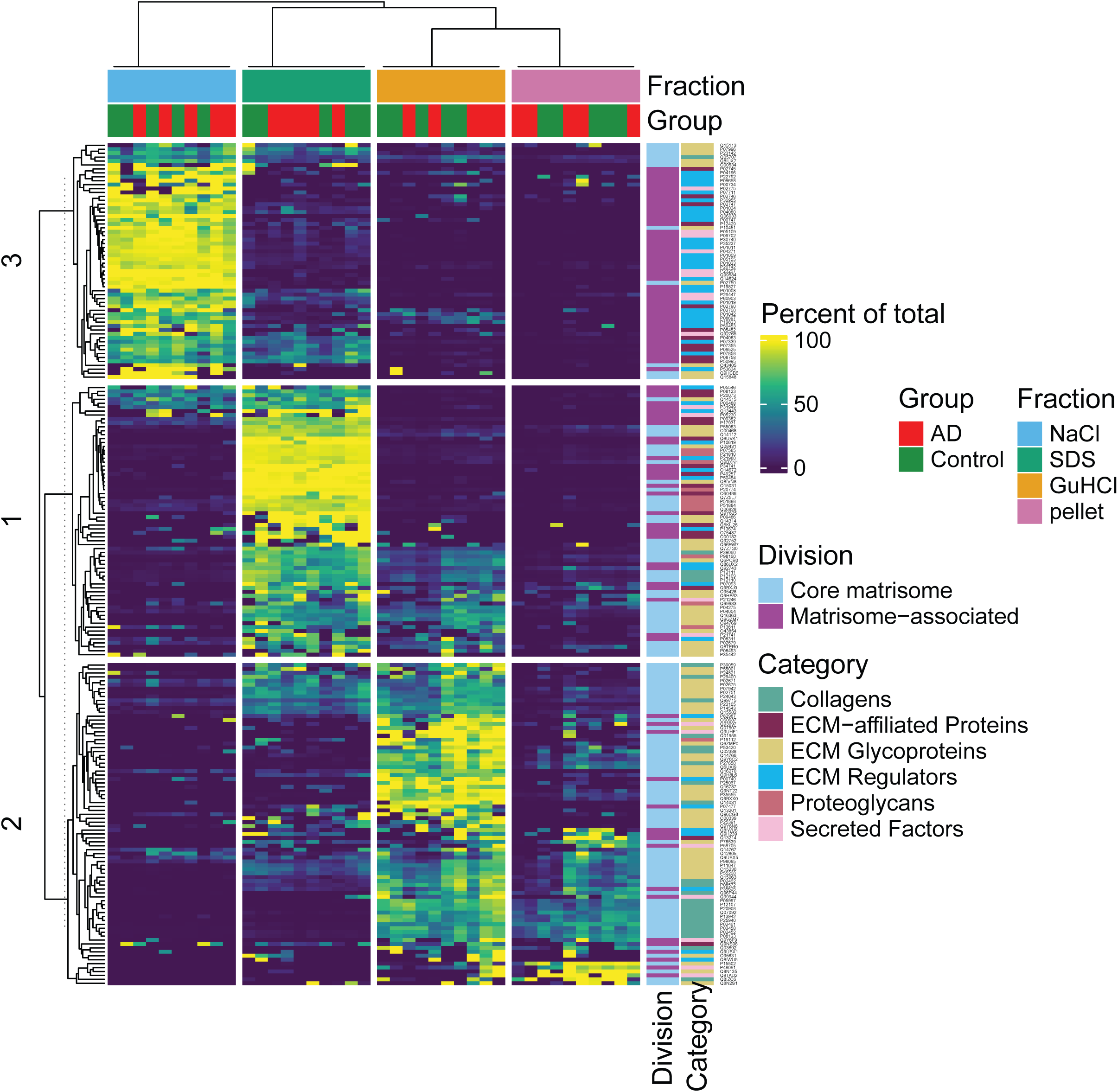
Biochemical fraction clustering of the human leptomeningeal matrisome. Heatmap shows normalized relative protein abundance across sequential biochemical fractions from human leptomeningeal samples derived from AD and CN individuals. Columns represent individual samples grouped by extraction fraction, including NaCl, SDS, GuHCl, and insoluble pellet, with fraction identity indicated by the top color bar. Disease status (AD vs. CN) for each sample is shown in the adjacent annotation bar. Columns are hierarchically clustered based on global proteomic similarity, illustrating fraction-driven rather than disease-driven separation of samples. Rows represent individual proteins identified across fractions and were hierarchically clustered using similarity in abundance patterns across all samples. The resulting rows were cut into three major clusters (labeled 1–3 along the left margin), each corresponding to a distinct biochemical solubility and extraction profile. These cluster labels are numerical identifiers without an implied ranking and serve to demarcate protein groups with shared fraction-enrichment patterns. White horizontal lines indicate the boundaries between these row clusters. Color intensity within the heatmap reflects normalized protein abundance, with higher values (yellow) indicating greater relative abundance and lower values (purple) indicating reduced or absent signal. The right-hand annotation panels indicate matrisome division (core matrisome vs. matrisome-associated) and functional matrisome category (collagens, ECM glycoproteins, proteoglycans, ECM regulators, ECM-affiliated proteins, and secreted factors).

**Table 3.**
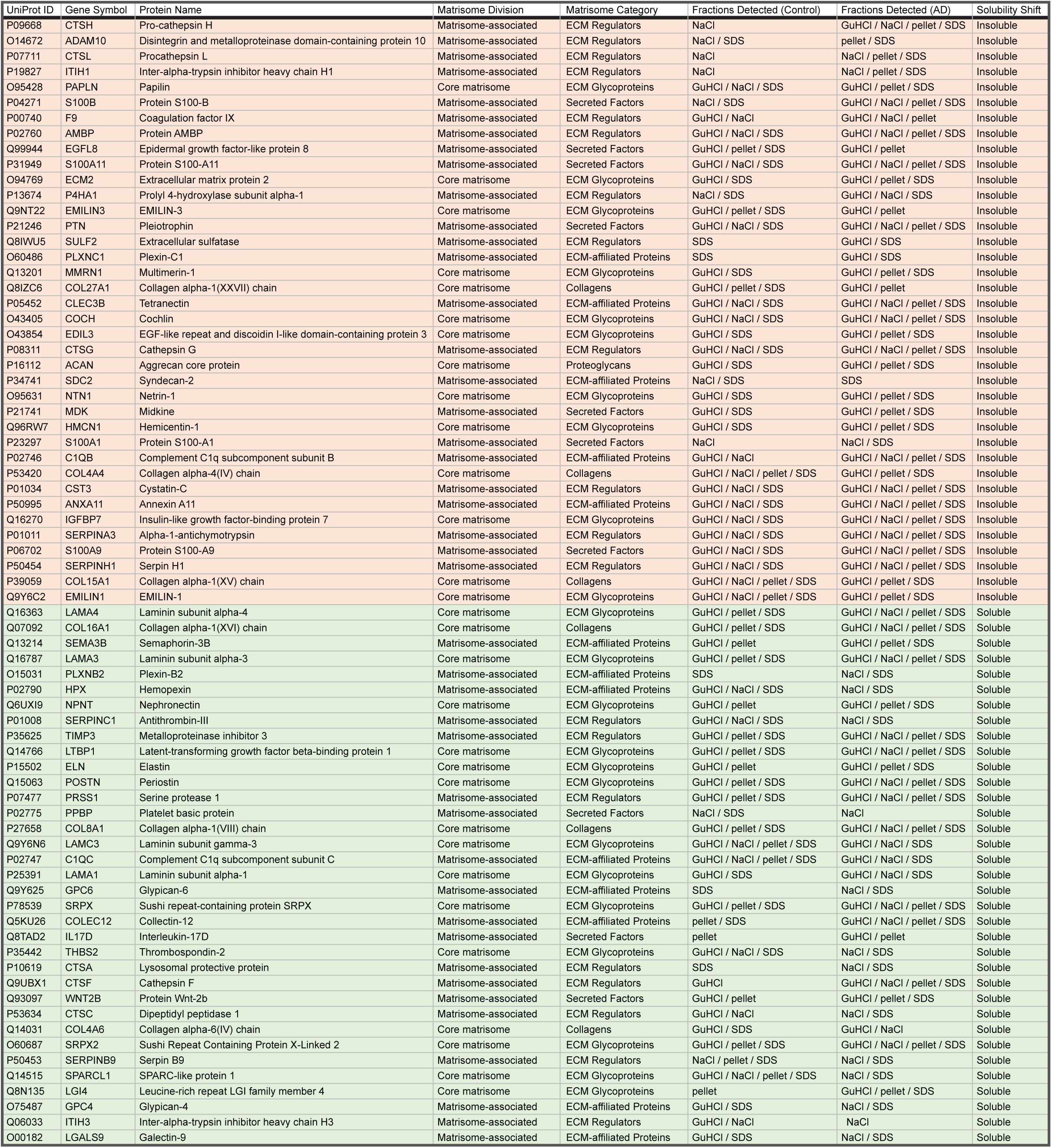
Protein solubility shifts based on the fraction distribution. Columns show gene symbol, UniProt ID, protein name, matrisome category and division, fractions where the protein was detected in CN and AD groups and solubility shift between the groups based on the enriched fractions in the experimental groups.

## DISCUSSION

Current evidence supports a connection between the leptomeninges and AD pathophysiology, including amyloid accumulation in early disease stages and a possible role in the clearance of AD biomarkers^8,68–70^. Original purification of Aβ included leptomeningeal tissue^71^. Neuropathological studies showed that Aβ deposition occurs in leptomeningeal vessels in CAA with later studies suggesting that impaired drainage at the meninges promotes vascular amyloid accumulation^71–73^. Recent analyses of CAA progression suggest that vascular deposition in cortical and leptomeningeal vessels represents an early component of disease pathology, possibly linked to disrupted clearance pathways^74,75^. These findings position the leptomeninges as a critical site for early processes in the development of AD pathology. In this study, we used a new proteomic pipeline to characterize changes in the ECM in AD in human leptomeninges as a discrete anatomical unit.

Protein composition of mouse meninges was investigated using genomic and proteomic approaches^3,6,76,77^. However, a significant limitation of rodent studies is that meningeal layers, leptomeninges and dura, cannot be clearly separated for individual analyses. Therefore, available datasets most likely represent the collective profile of all meningeal layers. It is challenging to pinpoint molecular changes specific to individual meningeal layers, which are distinct in their anatomical, structural, and physiological characteristics. Dura is a collagen-rich, rigid, and vascularized scaffold, and leptomeninges are mostly avascular, fragile, and thin tissues^2,10,19,24^. Species-specific differences are also significant. Human meninges are anatomically very different from rodent meninges. In humans, dura is an exceptionally tough, thick, and opaque layer, in rodents, it is fragile and translucent, pointing to the significant difference in structural ECM organization. In addition to anatomical considerations, our knowledge about remodeling of leptomeninges specifically in human disease remains very limited. Therefore, the use of human tissue in this work was critical for defining the matrisome in health and disease, and this is the first study to annotate leptomeningeal ECM.

Specimen preparation is a critical factor in proteomic analysis of the ECM. The solubility of the extracellular matrix varies widely depending on the specific proteins and the tissue context. Method comparisons for the matrisome analysis explicitly note that the insolubility of many ECM proteins and extraction protocols shape the coverage and recovery of ECM protein types^46,78^. Most proteoglycans and glycoproteins, some collagen types such as collagen IV, and fragments of ECM proteins generated by enzymatic degradation are soluble^30,79^. Collagens I, II, and III, crucial for structural integrity, are typically highly insoluble due to their cross-linking, as are elastin, laminin, and certain proteoglycans^46,80^. ECM components that have undergone extensive cross-linking, either through enzymatic or non-enzymatic means, become increasingly insoluble with age and tissue damage^81,82^. ECM regulators, secreted factors, and a subset of basement-membrane-associated proteins can be recovered under milder extraction conditions, whereas insoluble components, particularly core ECM, require guanidine or remain incompletely solubilized and underrepresented in one-step workflows or milder extraction conditions^46,78,79,83^. Therefore, to ensure complete ECM coverage in our datasets, we used a sequential, solubility-resolved extraction strategy that partitions proteins across NaCl-soluble, detergent-soluble (SDS), guanidine-soluble (GuHCl), and insoluble pellet fractions prior to digestion. The fractionation of our samples resulted in distinct proteome profiles across fractions and revealed a spectrum of leptomeningeal ECM biochemical accessibility.

Annotation of the matrisome proteins identified a glycoprotein-rich ECM dominated by basement membrane and perivascular ECM components. Multiple laminin heterotrimers were detected as well as perivascular ECM-associated proteins, including nephronectin (NPNT), tubulointerstitial nephritis antigen-like protein (TINAGL1), extracellular matrix protein 2 (ECM2), and hemicentin-1 (HMCN1). Elastic fiber and microfibrillar networks are prominent features of the leptomeningeal ECM. Together, these proteins define an ECM network consistent with tissues exposed to fluids and suggest that leptomeningeal ECM organization supports vascular elasticity, consistent with the anatomical association between blood vessels and leptomeninges. Multiple ECM regulators associated with transforming growth factor-β (TGFβ) signaling were identified within the leptomeningeal matrisome. These included latent TGFβ-binding proteins (LTBP1, LTBP2, LTBP4), TGFβ-induced protein (TGFBI), periostin (POSTN), adipocyte enhancer-binding protein 1 (AEBP1), procollagen C-endopeptidase enhancer 1 (PCOLCE), and collagen triple helix repeat-containing protein 1 (CTHRC1). The presence of these proteins indicates that the leptomeninges are capable of regulating ECM turnover and growth factor bioavailability. Proteins involved in organizing ECM junctions and barrier-like structures were prominent within the leptomeningeal matrisome. Laminins, nidogens, agrin, hemicentin-1, dermatopontin (DPT), and ECM2 together form a scaffold that anchors cellular elements and regulates ECM continuity across vascular and CSF-facing surfaces. The presence of these junctional components supports a role for the leptomeningeal ECM in maintaining boundary integrity in the CNS and regulating solute movement.

We identified three ECM proteins and one integrin as differentially expressed in AD based on the diagnosis alone. While integrins are not classified within the core matrisome framework used to annotate ECM datasets, they are membrane-spanning receptors that mediate bidirectional cell-matrix adhesion and signaling through binding to ECM ligands, including fibronectin, vitronectin, laminin, and collagen^30,52^. The identified ECM proteins, sulfatase 2 (SULF2), secreted frizzled-related protein 3 (sFRP3), and antithrombin-III (SERPINC1), belong to the groups of ECM regulators and secreted factors, pointing toward altered cell-matrix signaling and response rather than structural alterations in the leptomeninges in AD. The fourth protein, integrin beta-5 (ITGB5), combines with αV to form the αvβ5 receptor. This receptor binds fibronectin via two primary docking sites (the RGD motif in the tenth fibronectin type III domain and the ninth domain) and plays a role in ECM adhesion and outside-in signaling at the cell-matrix interface^84,85^.

SULF2 is an extracellular protein that modifies heparan sulfate (HS) sugar chains by selectively removing 6-O-sulfate groups from intact HS chains, altering the composition of cell-surface and matrix-associated HS Proteoglycans (HSPGs)^86^. By changing the sulfation pattern of the ECM, SULF2 regulates the binding and availability of HS-dependent growth factor families, including Wnt, FGF, and BMP^86,87^. SULF2 expression is decreased in the hippocampus and frontal lobe of AD patients compared with cognitively normal controls, suggesting that altered SULF2-mediated HS remodeling contributes to disease-related changes in neuronal cell signaling^88^. Highly sulfated HS subdomains removed by SULF-2 accumulate in cerebral Aβ plaques in both transgenic AD mouse models and humans in the entorhinal cortex and hippocampus. These HS subdomains are normally enriched in the vascular basement membrane of control tissues, but their distribution is altered in AD^89^. A multi-omic study of leptomeningeal tissue from CAA cases confirmed enrichment of HSPGs at sites of vascular Aβ deposition and found that Aβ40 abundance positively correlates with specific altered HS structures, including unsubstituted glucosamine residues^8^. That same study identified FRZB, a protein found significantly altered in AD in our specimens, as enriched in CAA-affected leptomeningeal vessels^8^. It is consistent with the pathology of our specimens, all of which were CAA-positive. CAA is a common feature of AD, present at autopsy in a substantial proportion of patients across multiple cohorts, with estimates of any-degree CAA in AD ranging from 80 to 90%^75,90^. sFRP3/FRZB has also been detected in human CSF by mass spectrometry, including peptide-level identification in CSF peptidomics datasets and increased in the serum of AD patients and in AD mouse models^91,92^. sFRP3/FRZB is an antagonist and extracellular modulator of the Wnt signaling pathway, which has been shown to be affected in AD^93,94^. It is possible that sFRP3 is specific to CAA rather than AD because these two factors are confounded in our study, however, these pathologies are closely linked^75,90^.

Antithrombin III (SERPINC1) is the major endogenous inhibitor of the coagulation cascade and a part of the fibrin coagulation regulatory system^95^. Reduced SERPINC1 levels in AD are consistent with the enrichment of fibrin-clot formation pathways identified in our Reactome analysis, suggesting diminished anticoagulant regulation and potentially increased thrombin activity, fibrin polymerization, and deposition within leptomeningeal vessels. Integrin beta-5 (ITGB5) encodes the β5 subunit of the αvβ5 integrin heterodimer, a vitronectin adhesion receptor, αvβ5 participates in ECM adhesion, endodermal cell differentiation, and TGF-signaling at the cell-matrix interface, consistent with TGF-β receptor signaling pathways^84,85^. In our dataset, TGF-β receptor signaling is one of the most significantly enriched pathways in the Reactome analysis of leptomeningeal ECM proteins. Our data align with previous studies that identified the upstream signaling axis (TGF-β), which appears to be present across meningeal compartments^6,77,96^.

Two proteins were identified only in CN leptomeninges under our inclusion criteria, i.e., present in a minimum of 3 samples per group: latent transforming growth factor-β binding protein 4 (LTBP4) and tenascin-R (TN-R). Both proteins are known regulators of ECM organization. LTBP4 is part of the latent transforming growth factor-β binding protein family^97,98^. LTBP4 binds and holds latent TGF-β1 within ECM components such as fibrillin and fibronectin^97,98^. This creates an extracellular reservoir of latent TGF-β that can later be activated through mechanical forces or proteolytic cleavage. In addition to regulating TGF-β bioavailability, LTBP4 contributes to elastic fiber and microfibril organization in the ECM, helping to control its activation and presentation to cell-surface receptors^98,99^. In this context, the fact that LTBP4 was not consistently detected in AD samples could indicate impaired TGF-β withholding and signaling within the meningeal ECM linked to control of fibroblast activity, matrix turnover, and tissue remodeling in AD. TN-R is an ECM glycoprotein that contributes to the assembly, stabilization, and organization of extracellular networks^100,101^. TN-R promotes the formation of aggrecan-rich ECM structures by clustering chondroitin sulfate proteoglycans within perineuronal nets^102,103^. Within the leptomeninges, low presence of TN-R could similarly destabilize ECM organization and impair the retention or localization of matrix-associated signaling molecules, including the TGF-β regulatory proteins. The proteins identified only in AD samples in our study are all secreted or ECM-associated factors, not structural core ECM components. This places them in the matrisome-associated category. These proteins belong to pathways regulating extracellular signaling and vascular remodeling, suggesting that AD-associated leptomeningeal changes in signaling environment rather than structural alterations.

Our fractionation approach revealed differences in ECM solubility between experimental groups. A group of soluble proteins that belong to ECM regulators, secreted factors, or ECM-affiliated proteins were newly enriched in more insoluble fractions in AD. Many of the proteins that control matrix organization and signaling are regulated by being held in tissue to varying degrees, some are more loosely associated, while others are tightly bound or crosslinked into the matrix^28–30^. This shift into insoluble fractions in AD may indicate that ECM regulators become more tightly matrix-bound, and secreted factors are increasingly sequestered within the insoluble ECM. Increased solubility of core matrisome proteins such as laminins, elastic fiber-associated proteins, and other structural extracellular matrix components may reflect disruption of higher-order matrix assemblies and reduced cross-linking. Laminins normally form tightly organized basement membrane scaffolds through interactions with collagen IV, nidogens, agrin, and integrins, and their increased extraction in soluble fractions may indicate basement membrane destabilization^23,25,104^. Similarly, elastic fiber-associated proteins, including elastin, fibulins, fibrillins, and EMILIN family members, are incorporated into highly cross-linked microfibrillar structures with low biochemical solubility. Increased solubility of these proteins is consistent with elastic fiber disassembly and impaired crosslinking^47,105^. Based on this, we propose that ECM remodeling in human leptomeninges occurs via biochemical accessibility and destabilization of core matrisome structural components.

This study has several limitations. Postmortem human tissue has inherent variability and comorbid pathologies, including CAA, which may confound disease-associated differences. In addition, the cohort size and type did not permit assessment of sex- and age-specific effects. In the future, more studies are required to functionally validate the identified pathways and proteins and establish their roles in AD. Overall, our study provides a robust workflow for identification and analysis of the meningeal ECM using mass spectrometry. We annotated and characterized the leptomeningeal matrisome in humans. Our findings demonstrated AD-associated shift of leptomeningeal ECM proteins’ solubility, affecting structural components, regulators, and signaling molecules. We identified ECM proteins that are differentially expressed in AD leptomeninges. These proteins belong to the associated matrisome, and we did not identify changes in expression among structural ECM components. Future studies and longitudinal analyses in human and animal models will determine how leptomeningeal ECM remodeling contributes to aging and AD pathophysiology.

## SUPPLEMENTARY MATERIALS

**Supplementary Table 1**. **List of proteins identified in human leptomeninges.** The proteins are identified by the protein ID number (column A) and the name (column B). Column C shows the matrisome status (matrisome or non-matrisome group).

**Supplementary Table 2. List of ECM proteins identified in human leptomeninges.** The proteins are organized by the division (core matrisome, associated matrisome and integrins) and by ECM type (collagens, glycoproteins, proteoglycans, secreted factors, ECM regulators and ECM-affiliated proteins). UniProt ID is displayed on the right.

**Supplementary Table 3. Distribution of leptomeningeal ECM proteins across fractions.** The table shows UniProt ID, gene symbol, protein name, matrisome division, matrisome category, fraction and solubility shift. Proteins were included and considered present if detected in ≥3 of 5 biological replicates within a given extraction fraction.

## DATA AVAILABILITY

The mass spectrometry raw proteomics data have been deposited to the ProteomeXchange Consortium via the PRIDE partner repository^106–108^ with the dataset identifier PXD077891 and 10.6019/PXD077891.

## ACKNOWLEDGEMENTS

We thank The Brain and Body Donation Program at the Banner Sun Health Research Institute for the provision of human biological materials. We thank Dr. George Chlipala (Research Informatics Core), University of Illinois at Chicago (UIC), who assisted with bioinformatics analysis.

