## Supplementary Table 3 for "Proteomic analysis of human leptomeningeal matrisome identifies changes in Alzheimer’s disease"

| UniProt ID | Gene Symbol | Protein Name | Matrisome Division | Matrisome Category | Fractions Detected (Control) | Fractions Detected (AD) | Solubility Shift |
|---|---|---|---|---|---|---|---|
| P09668 | CTSH | Pro-cathepsin H | Matrisome-associated | ECM Regulators | NaCl | GuHCl / NaCl / pellet / SDS | Insoluble |
| O14672 | ADAM10 | Disintegrin and metalloproteinase domain-containing protein 10 | Matrisome-associated | ECM Regulators | NaCl / SDS | pellet / SDS | Insoluble |
| P07711 | CTSL | Procathepsin L | Matrisome-associated | ECM Regulators | NaCl | NaCl / pellet / SDS | Insoluble |
| P19827 | ITIH1 | Inter-alpha-trypsin inhibitor heavy chain H1 | Matrisome-associated | ECM Regulators | NaCl | NaCl / pellet / SDS | Insoluble |
| O95428 | PAPLN | Papilin | Core matrisome | ECM Glycoproteins | GuHCl / NaCl / SDS | GuHCl / pellet / SDS | Insoluble |
| P04271 | S100B | Protein S100-B | Matrisome-associated | Secreted Factors | NaCl / SDS | GuHCl / NaCl / pellet / SDS | Insoluble |
| P00740 | F9 | Coagulation factor IX | Matrisome-associated | ECM Regulators | GuHCl / NaCl | GuHCl / NaCl / pellet | Insoluble |
| P02760 | AMBP | Protein AMBP | Matrisome-associated | ECM Regulators | GuHCl / NaCl / SDS | GuHCl / NaCl / pellet / SDS | Insoluble |
| Q99944 | EGFL8 | Epidermal growth factor-like protein 8 | Matrisome-associated | Secreted Factors | GuHCl / pellet / SDS | GuHCl / pellet | Insoluble |
| P31949 | S100A11 | Protein S100-A11 | Matrisome-associated | Secreted Factors | GuHCl / NaCl / SDS | GuHCl / NaCl / pellet / SDS | Insoluble |
| O94769 | ECM2 | Extracellular matrix protein 2 | Core matrisome | ECM Glycoproteins | GuHCl / SDS | GuHCl / pellet / SDS | Insoluble |
| P13674 | P4HA1 | Prolyl 4-hydroxylase subunit alpha-1 | Matrisome-associated | ECM Regulators | NaCl / SDS | GuHCl / NaCl / SDS | Insoluble |
| Q9NT22 | EMILIN3 | EMILIN-3 | Core matrisome | ECM Glycoproteins | GuHCl / pellet / SDS | GuHCl / pellet | Insoluble |
| P21246 | PTN | Pleiotrophin | Matrisome-associated | Secreted Factors | GuHCl / NaCl / SDS | GuHCl / NaCl / pellet / SDS | Insoluble |
| Q8IWU5 | SULF2 | Extracellular sulfatase | Matrisome-associated | ECM Regulators | SDS | GuHCl / SDS | Insoluble |
| O60486 | PLXNC1 | Plexin-C1 | Matrisome-associated | ECM-affiliated Proteins | SDS | GuHCl / SDS | Insoluble |
| Q13201 | MMRN1 | Multimerin-1 | Core matrisome | ECM Glycoproteins | GuHCl / SDS | GuHCl / pellet / SDS | Insoluble |
| Q8IZC6 | COL27A1 | Collagen alpha-1(XXVII) chain | Core matrisome | Collagens | GuHCl / pellet / SDS | GuHCl / pellet | Insoluble |
| P05452 | CLEC3B | Tetranectin | Matrisome-associated | ECM-affiliated Proteins | GuHCl / NaCl / SDS | GuHCl / NaCl / pellet / SDS | Insoluble |
| O43405 | COCH | Cochlin | Core matrisome | ECM Glycoproteins | GuHCl / NaCl / SDS | GuHCl / NaCl / pellet / SDS | Insoluble |
| O43854 | EDIL3 | EGF-like repeat and discoidin I-like domain-containing protein 3 | Core matrisome | ECM Glycoproteins | GuHCl / SDS | GuHCl / pellet / SDS | Insoluble |
| P08311 | CTSG | Cathepsin G | Matrisome-associated | ECM Regulators | GuHCl / NaCl / SDS | GuHCl / NaCl / pellet / SDS | Insoluble |
| P16112 | ACAN | Aggrecan core protein | Core matrisome | Proteoglycans | GuHCl / SDS | GuHCl / pellet / SDS | Insoluble |
| P34741 | SDC2 | Syndecan-2 | Matrisome-associated | ECM-affiliated Proteins | NaCl / SDS | SDS | Insoluble |
| O95631 | NTN1 | Netrin-1 | Core matrisome | ECM Glycoproteins | GuHCl / SDS | GuHCl / pellet / SDS | Insoluble |
| P21741 | MDK | Midkine | Matrisome-associated | Secreted Factors | GuHCl / SDS | GuHCl / pellet / SDS | Insoluble |
| Q96RW7 | HMCN1 | Hemicentin-1 | Core matrisome | ECM Glycoproteins | GuHCl / SDS | GuHCl / pellet / SDS | Insoluble |
| P23297 | S100A1 | Protein S100-A1 | Matrisome-associated | Secreted Factors | NaCl | NaCl / SDS | Insoluble |
| P02746 | C1QB | Complement C1q subcomponent subunit B | Matrisome-associated | ECM-affiliated Proteins | GuHCl / NaCl | GuHCl / NaCl / pellet / SDS | Insoluble |
| P53420 | COL4A4 | Collagen alpha-4(IV) chain | Core matrisome | Collagens | GuHCl / NaCl / pellet / SDS | GuHCl / pellet / SDS | Insoluble |
| P01034 | CST3 | Cystatin-C | Matrisome-associated | ECM Regulators | GuHCl / NaCl / SDS | GuHCl / NaCl / pellet / SDS | Insoluble |
| P50995 | ANXA11 | Annexin A11 | Matrisome-associated | ECM-affiliated Proteins | GuHCl / NaCl / SDS | GuHCl / NaCl / pellet / SDS | Insoluble |
| Q16270 | IGFBP7 | Insulin-like growth factor-binding protein 7 | Core matrisome | ECM Glycoproteins | GuHCl / NaCl / SDS | GuHCl / NaCl / pellet / SDS | Insoluble |
| P01011 | SERPINA3 | Alpha-1-antichymotrypsin | Matrisome-associated | ECM Regulators | GuHCl / NaCl / SDS | GuHCl / NaCl / pellet / SDS | Insoluble |
| P06702 | S100A9 | Protein S100-A9 | Matrisome-associated | Secreted Factors | GuHCl / NaCl / SDS | GuHCl / NaCl / pellet / SDS | Insoluble |
| P50454 | SERPINH1 | Serpin H1 | Matrisome-associated | ECM Regulators | GuHCl / NaCl / SDS | GuHCl / NaCl / pellet / SDS | Insoluble |
| P39059 | COL15A1 | Collagen alpha-1(XV) chain | Core matrisome | Collagens | GuHCl / NaCl / pellet / SDS | GuHCl / pellet / SDS | Insoluble |
| Q9Y6C2 | EMILIN1 | EMILIN-1 | Core matrisome | ECM Glycoproteins | GuHCl / NaCl / pellet / SDS | GuHCl / pellet / SDS | Insoluble |
| Q16363 | LAMA4 | Laminin subunit alpha-4 | Core matrisome | ECM Glycoproteins | GuHCl / pellet / SDS | GuHCl / NaCl / pellet / SDS | Soluble |
| Q07092 | COL16A1 | Collagen alpha-1(XVI) chain | Core matrisome | Collagens | GuHCl / pellet / SDS | GuHCl / NaCl / pellet / SDS | Soluble |
| Q13214 | SEMA3B | Semaphorin-3B | Matrisome-associated | ECM-affiliated Proteins | GuHCl / pellet | GuHCl / pellet / SDS | Soluble |
| Q16787 | LAMA3 | Laminin subunit alpha-3 | Core matrisome | ECM Glycoproteins | GuHCl / pellet / SDS | GuHCl / NaCl / pellet / SDS | Soluble |
| O15031 | PLXNB2 | Plexin-B2 | Matrisome-associated | ECM-affiliated Proteins | SDS | NaCl / SDS | Soluble |
| P02790 | HPX | Hemopexin | Matrisome-associated | ECM-affiliated Proteins | GuHCl / NaCl / SDS | NaCl / SDS | Soluble |
| Q6UXI9 | NPNT | Nephronectin | Core matrisome | ECM Glycoproteins | GuHCl / pellet | GuHCl / pellet / SDS | Soluble |
| P01008 | SERPINC1 | Antithrombin-III | Matrisome-associated | ECM Regulators | GuHCl / NaCl / SDS | NaCl / SDS | Soluble |
| P35625 | TIMP3 | Metalloproteinase inhibitor 3 | Matrisome-associated | ECM Regulators | GuHCl / pellet / SDS | GuHCl / NaCl / pellet / SDS | Soluble |
| Q14766 | LTBP1 | Latent-transforming growth factor beta-binding protein 1 | Core matrisome | ECM Glycoproteins | GuHCl / pellet / SDS | GuHCl / NaCl / pellet / SDS | Soluble |
| P15502 | ELN | Elastin | Core matrisome | ECM Glycoproteins | GuHCl / pellet | GuHCl / pellet / SDS | Soluble |
| Q15063 | POSTN | Periostin | Core matrisome | ECM Glycoproteins | GuHCl / pellet / SDS | GuHCl / NaCl / pellet / SDS | Soluble |
| P07477 | PRSS1 | Serine protease 1 | Matrisome-associated | ECM Regulators | GuHCl / pellet / SDS | GuHCl / NaCl / pellet / SDS | Soluble |
| P02775 | PPBP | Platelet basic protein | Matrisome-associated | Secreted Factors | NaCl / SDS | NaCl | Soluble |
| P27658 | COL8A1 | Collagen alpha-1(VIII) chain | Core matrisome | Collagens | GuHCl / pellet / SDS | GuHCl / NaCl / pellet / SDS | Soluble |
| Q9Y6N6 | LAMC3 | Laminin subunit gamma-3 | Core matrisome | ECM Glycoproteins | GuHCl / NaCl / pellet / SDS | GuHCl / NaCl / SDS | Soluble |
| P02747 | C1QC | Complement C1q subcomponent subunit C | Matrisome-associated | ECM-affiliated Proteins | GuHCl / NaCl / pellet / SDS | GuHCl / NaCl / SDS | Soluble |
| P25391 | LAMA1 | Laminin subunit alpha-1 | Core matrisome | ECM Glycoproteins | GuHCl / SDS | GuHCl / NaCl / SDS | Soluble |
| Q9Y625 | GPC6 | Glypican-6 | Matrisome-associated | ECM-affiliated Proteins | SDS | NaCl / SDS | Soluble |
| P78539 | SRPX | Sushi repeat-containing protein SRPX | Core matrisome | ECM Glycoproteins | GuHCl / pellet / SDS | GuHCl / NaCl / pellet / SDS | Soluble |
| Q5KU26 | COLEC12 | Collectin-12 | Matrisome-associated | ECM-affiliated Proteins | pellet / SDS | GuHCl / NaCl / pellet / SDS | Soluble |
| Q8TAD2 | IL17D | Interleukin-17D | Matrisome-associated | Secreted Factors | pellet | GuHCl / pellet | Soluble |
| P35442 | THBS2 | Thrombospondin-2 | Core matrisome | ECM Glycoproteins | GuHCl / NaCl / SDS | NaCl / SDS | Soluble |
| P10619 | CTSA | Lysosomal protective protein | Matrisome-associated | ECM Regulators | SDS | NaCl / SDS | Soluble |
| Q9UBX1 | CTSF | Cathepsin F | Matrisome-associated | ECM Regulators | GuHCl | GuHCl / NaCl / pellet / SDS | Soluble |
| Q93097 | WNT2B | Protein Wnt-2b | Matrisome-associated | Secreted Factors | GuHCl / pellet | GuHCl / pellet / SDS | Soluble |
| P53634 | CTSC | Dipeptidyl peptidase 1 | Matrisome-associated | ECM Regulators | GuHCl / NaCl | NaCl / SDS | Soluble |
| Q14031 | COL4A6 | Collagen alpha-6(IV) chain | Core matrisome | Collagens | GuHCl / SDS | GuHCl / NaCl | Soluble |
| O60687 | SRPX2 | Sushi Repeat Containing Protein X-Linked 2 | Core matrisome | ECM Glycoproteins | GuHCl / pellet / SDS | GuHCl / NaCl / pellet / SDS | Soluble |
| P50453 | SERPINB9 | Serpin B9 | Matrisome-associated | ECM Regulators | NaCl / pellet / SDS | NaCl / SDS | Soluble |
| Q14515 | SPARCL1 | SPARC-like protein 1 | Core matrisome | ECM Glycoproteins | GuHCl / NaCl / pellet / SDS | NaCl / SDS | Soluble |
| Q8N135 | LGI4 | Leucine-rich repeat LGI family member 4 | Core matrisome | ECM Glycoproteins | pellet | GuHCl / pellet / SDS | Soluble |
| O75487 | GPC4 | Glypican-4 | Matrisome-associated | ECM-affiliated Proteins | GuHCl / SDS | NaCl / SDS | Soluble |
| Q06033 | ITIH3 | Inter-alpha-trypsin inhibitor heavy chain H3 | Matrisome-associated | ECM Regulators | GuHCl / NaCl | NaCl | Soluble |
| O00182 | LGALS9 | Galectin-9 | Matrisome-associated | ECM-affiliated Proteins | GuHCl / SDS | NaCl / SDS | Soluble |
