## Supplementary Table 2 for "Proteomic analysis of human leptomeningeal matrisome identifies changes in Alzheimer’s disease"

| Division | Category | Proetin Name | UniprotID |
|---|---|---|---|
| Core Matrisome | Collagens | Collagen alpha-2(V) chain | P05997 |
| | | Collagen alpha-2(VI) chain | P12110 |
| | | Collagen alpha-3(VI) chain | P12111 |
| | | Collagen alpha-1(XIV) chain | Q05707 |
| | | Collagen alpha-1(XVI) chain | Q07092 |
| | | Collagen alpha-6(IV) chain | Q14031 |
| | | Collagen alpha-1(XII) chain | Q99715 |
| | | Collagen alpha-1(IV) chain | P02462 |
| | | Collagen alpha-2(XI) chain | P13942 |
| | | Collagen alpha-1(V) chain | P20908 |
| | | Collagen alpha-1(XXVII) chain | Q8IZC6 |
| | | Collagen alpha-1(XV) chain | P39059 |
| | | Collagen alpha-2(I) chain | P08123 |
| | | Collagen alpha-1(II) chain | P02458 |
| | | Collagen alpha-1(VII) chain | Q02388 |
| | | Collagen alpha-2(IV) chain | P08572 |
| | | Collagen alpha-1(VI) chain | P12109 |
| | | Collagen alpha-1(XVIII) chain | P39060 |
| | | Collagen alpha-1(III) chain | P02461 |
| | | Collagen alpha-1(VIII) chain | P27658 |
| | | Collagen alpha-1(I) chain | P02452 |
| | | Collagen alpha-5(IV) chain | P29400 |
| | | Collagen alpha-4(IV) chain | P53420 |
| | | Collagen alpha-1(XXI) chain | Q96P44 |
| | | Collagen alpha-1(XI) chain | P12107 |
| | | Collagen alpha-2(VIII) chain | P25067 |
| | | Collagen alpha-3(V) chain | P25940 |
| | | Collagen alpha-3(IV) chain | Q01955 |
| | | Collagen alpha-1(X) chain | Q03692 |
| | ECM Glycoproteins | Latent-transforming growth factor beta-binding protein 1 | Q14766 |
| | | Transforming growth factor-beta-induced protein ig-h3 | Q15582 |
| | | Matrix Gla protein | P08493 |
| | | Thrombospondin-2 | P35442 |
| | | Laminin subunit beta-2 | P55268 |
| | | Sushi repeat-containing protein SRPX | P78539 |
| | | Dermatopontin | Q07507 |
| | | Periostin | Q15063 |
| | | Insulin-like growth factor-binding protein 7 | Q16270 |
| | | Laminin subunit alpha-3 | Q16787 |
| | | Thrombospondin type-1 domain-containing protein 4 | Q6ZMP0 |
| | | Adipocyte enhancer-binding protein 1 | Q8IUX7 |
| | | Tubulointerstitial nephritis antigen-like | Q9GZM7 |
| | | Tenascin-X | P22105 |
| | | Fibrillin-1 | P35555 |
| | | Nidogen-1 | P14543 |
| | | Spondin-1 | Q9HCB6 |
| | | Laminin subunit gamma-1 | P11047 |
| | | von Willebrand factor A domain-containing protein 5A | O00534 |
| | | EGF-like repeat and discoidin I-like domain-containing protein 3 | O43854 |
| | | Vitronectin | P04004 |
| | | von Willebrand factor | P04275 |
| | | Elastin | P15502 |
| | | Multimerin-1 | Q13201 |
| | | Fibroleukin | Q14314 |
| | | von Willebrand factor A domain-containing protein 1 | Q6PCB0 |
| | | Nephronectin | Q6UXI9 |
| | | Leucine-rich repeat LGI family member 4 | Q8N135 |
| | | Tenascin-R | Q92752 |
| | | EMILIN-3 | Q9NT22 |
| | | Multimerin-2 | Q9H8L6 |
| | | Nidogen-2 | Q14112 |
| | | Laminin subunit alpha-1 | P25391 |
| | | Lactadherin | Q08431 |
| | | SPARC-like protein 1 | Q14515 |
| | | Target of Nesh-SH3 | Q7Z7G0 |
| | | SPARC | P09486 |
| | | Fibronectin | P02751 |
| | | Fibulin-1 | P23142 |
| | | Laminin subunit gamma-3 | Q9Y6N6 |
| | | Fibulin-2 | P98095 |
| | | Papilin | O95428 |
| | | Netrin-4 | Q9HB63 |
| | | Sushi repeat-containing protein SRPX2 | O60687 |
| | | Tenascin | P24821 |
| | | Sushi, nidogen and EGF-like domain-containing protein 1 | Q8TER0 |
| | | EGF-containing fibulin-like extracellular matrix protein 1 | Q12805 |
| | | Hemicentin-1 | Q96RW7 |
| | | EMILIN-1 | Q9Y6C2 |
| | | Collagen triple helix repeat-containing protein 1 | Q96CG8 |
| | | Microfibril-associated glycoprotein 4 | P55083 |
| | | Latent-transforming growth factor beta-binding protein 2 | Q14767 |
| | | Agrin | O00468 |
| | | Fibrinogen alpha chain | P02671 |
| | | Procollagen C-endopeptidase enhancer 1 | Q15113 |
| | | Thrombospondin-1 | P07996 |
| | | Laminin subunit alpha-5 | O15230 |
| | | Extracellular matrix protein 2 | O94769 |
| | | Fibrinogen beta chain | P02675 |
| | | Fibrinogen gamma chain | P02679 |
| | | Laminin subunit alpha-4 | Q16363 |
| | | Fibulin-5 | Q9UBX5 |
| | | Adiponectin | Q15848 |
| | | Laminin subunit beta-1 | P07942 |
| | | Cochlin | O43405 |
| | | Matrilin-2 | O00339 |
| | | Netrin-1 | O95631 |
| | | Leucine-rich alpha-2-glycoprotein | P02750 |
| | | Osteopontin | P10451 |
| | | Laminin subunit alpha-2 | P24043 |
| | | Microfibrillar-associated protein 2 | P55001 |
| | | Somatomedin-B and thrombospondin type-1 domain-containing protein | Q8IVN8 |
| | | Latent-transforming growth factor beta-binding protein 4 | Q8N2S1 |
| | | EMILIN-2 | Q9BXX0 |
| | Proteoglycans | Podocan | Q7Z5L7 |
| | | Versican core protein | P13611 |
| | | Aggrecan core protein | P16112 |
| | | Basement membrane-specific heparan sulfate proteoglycan core protein | P98160 |
| | | Asporin | Q9BXN1 |
| | | Osteomodulin | Q99983 |
| | | Fibromodulin | Q06828 |
| | | Lumican | P51884 |
| | | Mimecan | P20774 |
| | | Biglycan | P21810 |
| | | Decorin | P07585 |
| | | Prolargin | P51888 |
| Matrisome-Associated | Secreted Factors | Midkine | P21741 |
| | | Protein S100-A1 | P23297 |
| | | Protein S100-A4 | P26447 |
| | | Secreted frizzled-related protein 3 | Q92765 |
| | | Pleiotrophin | P21246 |
| | | Platelet basic protein | P02775 |
| | | Protein Wnt-2b | Q93097 |
| | | Epidermal growth factor-like protein 8 | Q99944 |
| | | Epidermal growth factor-like protein 7 | Q9UHF1 |
| | | Fibroblast growth factor 1 | P05230 |
| | | Protein S100-A9 | P06702 |
| | | Protein S100-A11 | P31949 |
| | | Protein S100-B | P04271 |
| | | Protein S100-A8 | P05109 |
| | | Stromal cell-derived factor 1 | P48061 |
| | | Protein Wnt-4 | P56705 |
| | | Protein S100-A10 | P60903 |
| | | Interleukin-17D | Q8TAD2 |
| | | Protein S100-A13 | Q99584 |
| | | Protein Wnt-6 | Q9Y6F9 |
| | ECM Regulators | Extracellular sulfatase Sulf-1 | Q8IWU6 |
| | | Serine protease 1 | P07477 |
| | | Disintegrin and metalloproteinase domain-containing protein 10 | O14672 |
| | | Coagulation factor XIII A chain | P00488 |
| | | Plasma protease C1 inhibitor | P05155 |
| | | Cathepsin B | P07858 |
| | | Protein-glutamine gamma-glutamyltransferase 2 | P21980 |
| | | Serpin B6 | P35237 |
| | | Metalloproteinase inhibitor 3 | P35625 |
| | | Cathepsin F | Q9UBX1 |
| | | Cathepsin G | P08311 |
| | | Dipeptidyl peptidase 1 | P53634 |
| | | Procathepsin L | P07711 |
| | | Alpha-2-macroglobulin | P01023 |
| | | Antithrombin-III | P01008 |
| | | Alpha-1-antichymotrypsin | P01011 |
| | | Angiotensinogen | P01019 |
| | | Pro-cathepsin H | P09668 |
| | | Inter-alpha-trypsin inhibitor heavy chain H1 | P19827 |
| | | Carboxypeptidase N subunit 2 | P22792 |
| | | Pigment epithelium-derived factor | P36955 |
| | | ADAMTS-like protein 3 | P82987 |
| | | Matrix metalloproteinase-28 | Q9H239 |
| | | Serpin B9 | P50453 |
| | | Heparin cofactor 2 | P05546 |
| | | Leukocyte elastase inhibitor | P30740 |
| | | Cystatin-C | P01034 |
| | | Cystatin-B | P04080 |
| | | Alpha-2-antiplasmin | P08697 |
| | | Inter-alpha-trypsin inhibitor heavy chain H4 | Q14624 |
| | | Prolyl 4-hydroxylase subunit alpha-1 | P13674 |
| | | Inter-alpha-trypsin inhibitor heavy chain H2 | P19823 |
| | | Histidine-rich glycoprotein | P04196 |
| | | Serine protease HTRA1 | Q92743 |
| | | Serpin H1 | P50454 |
| | | Alpha-1-antitrypsin | P01009 |
| | | Kininogen-1 | P01042 |
| | | Plasminogen | P00747 |
| | | Protein AMBP | P02760 |
| | | Lysosomal protective protein | P10619 |
| | | Pregnancy zone protein | P20742 |
| | | Inter-alpha-trypsin inhibitor heavy chain H5 | Q86UX2 |
| | | Cathepsin D | P07339 |
| | | Glia-derived nexin | P07093 |
| | | Disintegrin and metalloproteinase domain-containing protein 9 | Q13443 |
| | | Coagulation factor IX | P00740 |
| | | Inter-alpha-trypsin inhibitor heavy chain H3 | Q06033 |
| | | Prothrombin | P00734 |
| | | Extracellular sulfatase Sulf-2 | Q8IWU5 |
| | ECM-affiliated Proteins | Complement C1q subcomponent subunit B | P02746 |
| | | Semaphorin-3B | Q13214 |
| | | Annexin A4 | P09525 |
| | | Plexin-B2 | O15031 |
| | | Glypican-4 | O75487 |
| | | Complement C1q subcomponent subunit A | P02745 |
| | | Annexin A1 | P04083 |
| | | Tetranectin | P05452 |
| | | Annexin A2 | P07355 |
| | | Annexin A6 | P08133 |
| | | Annexin A5 | P08758 |
| | | Syndecan-2 | P34741 |
| | | Protein ERGIC-53 | P49257 |
| | | Annexin A11 | P50995 |
| | | Glypican-6 | Q9Y625 |
| | | Complement C1q subcomponent subunit C | P02747 |
| | | Plexin-C1 | O60486 |
| | | Hemopexin | P02790 |
| | | Chondroitin sulfate proteoglycan 4 | Q6UVK1 |
| | | Complement C1q tumor necrosis factor-related protein 5 | Q9BXJ0 |
| | | Semaphorin-3G | Q9NS98 |
| | | Collectin-12 | Q5KU26 |
| | | Annexin A3 | P12429 |
| | | Galectin-1 | P09382 |
| | | Galectin-3 | P17931 |
| | | Galectin-9 | O00182 |
| | | Annexin A7 | P20073 |
| Integrins | | Integrin alpha-1 | P56199 |
| | | Integrin alpha-3 | P26006 |
| | | Integrin alpha-5 | P08648 |
| | | Integrin alpha-6 | P23229 |
| | | Integrin alpha-7 | Q13683 |
| | | Integrin alpha-8 | P53708 |
| | | Integrin alpha-IIb | P08514 |
| | | Integrin alpha-M | P11215 |
| | | Integrin alpha-V | P06756 |
| | | Integrin beta-1 | P05556 |
| | | Integrin beta-2 | P05107 |
| | | Integrin beta-3 | P05106 |
| | | Integrin beta-4 | P16144 |
| | | Integrin beta-5 | P18084 |
| | | Integrin beta-6 | P18564 |
| | | Integrin-linked protein kinase | Q13418 |
